# Rapid and dynamic reprogramming within the tumor microenvironment drives EDA-CAR-T dysfunction and compromised therapeutic efficacy in solid tumors

**DOI:** 10.64898/2026.04.29.721801

**Authors:** Rodrigo Redondo-Frutos, Pedro Justicia-Lirio, Maria E. Cervantes-Calleja, Patxi San Martin-Uriz, Paula Aguirre-Ruiz, Lorea Jordana-Urriza, Maider Garnica-Suberviola, Sergio Camara-Peña, Diego Alignani, Aitziber Lopez, Saray Rodriguez-Diaz, Rebeca Martinez-Turrillas, Marta Gorraiz, Derya Bakirdöğen, Arianna Pocaterra, Susana Inogés, Ascensión López-Díaz de Cerio, Hanna Algül, Anna Mondino, Mikel Hernaez, Juan J. Lasarte, Felipe Prosper, Teresa Lozano, Juan R. Rodriguez-Madoz

**Author notes:** These authors share senior authorship (Teresa Lozano, Juan R. Rodriguez-Madoz). Correspondence: Juan R. Rodriguez-Madoz.

## Abstract

**Background:** Chimeric antigen receptor (CAR)-T cell therapies efficacy in solid tumors remains limited, largely due to the profoundly immunosuppressive tumor microenvironment (TME) which drives CAR-T cells to dysfunction and poor persistence. A comprehensive understanding of the dynamic interplay between CAR-T cells and the TME is therefore critical for the rational design of more effective CAR-T strategies for solid cancers.

**Methods:** Here, we performed single-cell RNA sequencing of tumor samples from immunocompetent mice treated with stroma-targeting EDA-CAR-T cells, profiling CAR-T cell states and TME programs at the peak of antitumor response and during subsequent tumor progression.

**Results:** Our analysis revealed a marked temporal remodeling of EDA-CAR-T cells within the TME, where early antitumor efficacy is associated with concurrent expansion of cytotoxic effector CD8⁺ CAR-T cells and activation of memory CD4⁺ CAR-T subsets. Moreover, EDA-CAR-T cells effectively engaged the myeloid compartment, resulting in strengthened communication networks involving T cell activation. However, by tumor progression, EDA-CAR-T cells suffered a widespread transcriptional reprogramming towards dysfunction, characterized by loss of effector programs alongside induction of exhaustion and immunoregulatory pathways within the TME, including PD-L1/PD-L2 and TGFβ signaling, which impairs sustained immune responses. Notably, early CAR-T cell activation led to increased susceptibility to TME-mediated immunosuppression, revealing EDA-CAR-T-specific soluble galectin-mediated cell-to-cell interaction networks.

**Conclusions:** Together, this works offers a high-resolution view of CAR-T cell dynamics within the solid TME, uncovering cellular and molecular mechanisms of rapid functional decline and identifying regulatory pathways within the TME that can be exploited to improve CAR-T cell therapy efficacy in solid tumors.

**KEY MESSAGES OF THE ARTICLE:** *What is already known on this topic:* The determinants of CAR-T cell therapeutic efficacy in solid tumors remain poorly defined, largely due to the complexity of the immunosuppressive tumor microenvironment. In this effort, it is necessary to perform comprehensive and detailed mechanistic studies that capture CAR-T cell dynamics within the solid tumor microenvironment to understand treatment failure.

*What this study adds:* We performed single-cell profiling of stroma-targeting EDA-CAR-T cells, revealing their dynamic reprogramming toward dysfunction within the solid tumor microenvironment. We dissected CAR-T cell states and their cell-to-cell interactions with the tumor microenvironment across response and tumor progression and identified mechanisms linking CAR-T cell functionality and therapeutic failure.

*How this study might affect research, practice or policy:* This study provides comprehensive mechanistic insights from an immunocompetent model that can be leveraged to identify shared determinants of CAR-T cell functionality in solid tumors and potentially guide the rational development of improved CAR-T cell therapies.

## INTRODUCTION

Chimeric antigen receptor (CAR)-T cell therapy has achieved remarkable clinical success in the treatment of hematological malignancies, leading to long-term remissions in B-cell cancers through targeting of CD19 and B-cell maturation antigen (BCMA)^1–3^. In contrast, efficacy in solid tumors remains limited, with clinical benefit largely restricted to small, heavily pretreated patients, and characterized by no durable responses and rapid disease progression^4–6^.

Beyond antigen recognition, solid tumors impose additional barriers, including restricted tissue infiltration and a highly immunosuppressive tumor microenvironment (TME)^7–9^. The TME represents a central determinant of CAR-T cell failure in solid tumors. It harbors a heterogeneous composition of malignant and non-malignant cells that integrate physical barriers with metabolic and immune suppressive cues. CAR-T cells interactions within this milieu restrict trafficking and intratumoral infiltration and persistence, promote T cell dysfunction and facilitate immune evasion, ultimately limiting sustained antitumor activity^1,6,10,11^. Therefore, dissecting CAR-T cell crosstalk in the TME is critical for rational therapeutic improvements.

Single-cell RNA sequencing (scRNA-seq) enables unbiased, high-resolution profiling to dissect cellular heterogeneity and functional states within the TME^12^. scRNA-seq studies has uncovered extensive diversity among tumor-infiltrating T lymphocytes (TILs), demonstrating that distinct T cell states contribute differently to tumor control and immunotherapy outcomes^13–16^. Applied to CAR-T cell therapies, scRNA-seq allows direct interrogation of CAR-T cell states within the complex tumor niche. To date, mechanistic insights into CAR-T biology derive largely from hematological malignancies. In B-cell cancers, scRNA-seq studies have defined transcriptional programs associated with CAR-T expansion and long-term persistence^17–23^. However, these paradigms are unlikely to extend to solid tumors, where antigen exposure is spatially restricted, peripheral expansion is limited, and T cell activation and maintained functionality must occur predominantly within the hostile TME^24^. Consequently, determinants of CAR-T efficacy in solid tumors remain poorly defined, highlighting the need for detailed mechanistic studies that capture CAR-T cell dynamics within the TME.

We have previously developed a CAR specific for the extra-domain A (EDA) splice variant of fibronectin (EDA-CAR), a stromal antigen broadly present across multiple tumor types. EDA-CAR-T cells demonstrated robust antitumor activity in immunocompetent models in which tumor cells overexpress EDA^25^. However, its therapeutic efficacy was significantly compromised in murine models where EDA is predominantly deposited within the extracellular matrix or localized to the tumor endothelial basement membrane^25^.

To better understand the mechanisms underlying this impairment in response, in this work we performed a comprehensive multi-omic analysis of EDA-CAR-T cell therapy in a challenging immunocompetent model, focusing on tumor-infiltrating CAR-T cells and the surrounding TME. Using single-cell RNA sequencing, we profiled the cellular composition, functional states and intercellular communication networks of CAR-T cells within the TME at both the peak of response and at the time of tumor progression. These integrative analyses identified key determinants of CAR-T cell activation, differentiation, and dysfunction within the TME, uncovering potential targets for therapeutic optimization, and providing a translational framework to guide the rational design of next-generation CAR-T strategies.

## MATERIALS AND METHODS

### Mice

Female 129Sv mice (10-12 weeks old) were obtained from Janvier Labs (Le Genest-Saint-Isle, France) and housed under specific pathogen-free conditions at the animal facility of the Cima Universidad de Navarra. All animal procedures were approved by the local ethical committee and performed in accordance with institutional guidelines.

### Cell culture

The murine teratocarcinoma F9 cell line (ATCC) was cultured in RPMI 1640 supplemented with GlutaMAX™, 10% fetal bovine serum (FBS), 1x penicillin-streptomycin, 2 mM L-glutamine, 10 mM HEPES buffer, 1x MEM non-essential amino acids, 1 mM sodium pyruvate, 50 µM β-mercaptoethanol and 2.5 µg/mL amphotericin B (hereafter referred to as complete medium). Platinum Ecotropic (ATCC) packaging cells were maintained in complete DMEM containing 100 µg/mL puromycin and 10 µg/mL blasticidin for selection. All cell lines were maintained at 37°C in 6.5% CO₂.

### CAR constructs and retroviral production

The murine EDA-CAR consisted of the anti-EDA F8 scFv fused to a murine 4-1BB-CD3ζ signaling cassette and linked via a F2A self-cleaving peptide sequence to eGFP. An irrelevant PSMA-CAR (anti-human PSMA scFv from mouse hybridoma J591) cloned in the same backbone served as control. Constructs were synthesized by GenScript. Retrovirus was produced by transfecting Plat-E cells with retroviral plasmid DNA and pCL-Eco packaging vector using Lipofectamine 2000. Viral supernatants were collected at 48 and 72 hours post-transfection and filtered (0.45 µm).

### Murine T cell isolation and retroviral transduction

CD4⁺ and CD8⁺ T cells were isolated from spleens and lymph nodes of 129Sv mice by negative magnetic selection (Miltenyi Biotec). Purified T cells were activated with anti-CD3/CD28 Dynabeads (1:2 bead-to-cell ratio) in complete RPMI supplemented with recombinant human IL-2 (rhIL-2, 100 IU/mL). Activated T cells were transduced with retroviral supernatant in the presence of protamine sulfate by spinoculation. Transduction was repeated 24 hours later. T cells were expanded in rhIL-2-containing complete RPMI medium and used on day 5. Transduction efficiency was assessed by GFP expression via flow cytometry.

### Tumor model and adoptive cell transfer

Mice were injected subcutaneously with 3×10⁶ F9 cells in the right flank. Five days later, animals received lymphodepleting total-body γ-irradiation (4 Gy), followed by intravenous infusion of 1×10⁷ CAR-T cells (1:1 CD4⁺:CD8⁺). Recombinant human rhIL-2 (20,000 IU) was administered intraperitoneally daily for four consecutive days.

### Tissue collection, processing and single-cell preparation

Tumors were excised, mechanically dissociated, and enzymatically digested with collagenase D and DNase I. Digestion was stopped with EDTA, and suspensions were filtered and subjected to red blood cell lysis. Spleens were mechanically dissociated through cell strainers to obtain single-cell suspensions, followed by red blood cell lysis. All samples were washed and resuspended in PBS.

### Flow cytometry

Single-cell suspensions were stained with fluorochrome-conjugated antibodies, and CAR-T cells were identified by GFP. The viability dye EMA was used to exclude dead cells. Data was acquired on a FACSCanto-II cytometer and analyzed with FlowJo software (v10.0.0). For scRNA-seq experiments, tumor suspensions were first incubated with Fc receptor-blocking antibody and then stained with TotalSeq™ hashing antibodies for sample multiplexing together with surface fluorochrome-conjugated antibodies. Viable CD45⁺, CD45^-^, and GFP⁺ CAR-T populations were sorted on a FACSAria™ IIU cell sorter. All fluorochrome-conjugated and TotalSeq™ hashing antibodies used are listed in Supplementary Methods.

### Single-cell RNA sequencing and TCR profiling

FACS-sorted populations were processed using the 10x Genomics Chromium platform for 5′ gene expression and paired TCR α/β sequencing (Table S1). Gene expression libraries were prepared using the Chromium Single Cell 5′ Reagent Kit v3. Libraries were sequenced on an Illumina NextSeq2000 at a depth of 30,000-50,000 reads per cell. TCR libraries were generated using the Chromium V(D)J Enrichment Kit for mouse T cells and sequenced at 7,000-15,000 reads per cell. Hashtag oligonucleotide libraries for sample multiplexing were prepared according to manufacturer instructions and sequenced at 5000-10000 reads per cell.

### Single-cell data processing and integration

Raw sequencing data were processed using Cell Ranger for alignment to the mouse reference genome (GRCm38), barcode assignment, and UMI counting. Downstream analyses were performed in R using Seurat. Ambient RNA contamination was corrected with SoupX, and doublets were identified using DoubletFinder and hashtag demultiplexing (HTODemux). Low-quality cells were excluded based on gene counts, UMI thresholds, mitochondrial and ribosomal content, and doublet classification. Data were log-normalized, and 2,000 highly variable genes were identified. Batch effects across experiments were corrected using Harmony.

### Cell type clustering and annotation

Dimensionality reduction was performed by PCA, and the top 20 principal components were used for Uniform Manifold Approximation and Projection (UMAP) visualization and Louvain clustering. TME cell populations were annotated based on canonical marker gene expression. T cells and myeloid populations were subsetted and reclustered for higher-resolution analysis. Cluster-specific marker genes were identified using differential expression testing (average log fold-change > 0.25; detected in >10% of cells). CAR-T cells were identified by detection of GFP transcripts. All marker genes used for cluster annotation are listed in the Supplementary Methods.

### TCR repertoire analysis

TCR clonotypes were analyzed using the scRepertoire package. Clonal expansion and diversity metrics were assessed across time points and treatment conditions.

### Pseudobulk differential expression and pathway analyses

Differential gene expression was performed using a pseudobulk strategy. UMI counts were aggregated per biological replicate and analyzed with DESeq2 using negative binomial generalized linear models. Likelihood ratio tests were used to assess treatment-dependent effects, with PSMA-CAR-T as reference. Genes with FDR < 0.05 were considered significant. Gene Ontology (GO) enrichment analysis was conducted using clusterProfiler (enrichGO function). Gene set enrichment analysis (GSEA) was performed on ranked gene lists using curated immune-related signatures. Pathways with FDR < 0.25 were considered significantly enriched. Gene set variation analysis (GSVA) was applied to pseudobulk expression matrices to quantify pathway activity differences across conditions. Statistical comparisons were performed using non-parametric tests with Benjamini-Hochberg correction.

### Cell-cell communication analysis

Intercellular communication networks were inferred using CellChat, integrating ligand-receptor interaction databases. Interaction probabilities were computed based on expression of ligand-receptor pairs across annotated cell types. Differential signaling strength between conditions was evaluated using Z-score-normalized interaction scores and permutation-based significance testing.

### Statistical analysis

Statistical analyses were performed in R and GraphPad Prism. The different tests used in this work are indicated in figure legends.

## RESULTS

### Comprehensive single-cell RNA-seq analysis of TME after treatment with EDA-CAR-T cells

We have previously shown that EDA-CAR-T cells presented strong antitumoral efficacy in several immunocompetent models overexpressing EDA. However, their efficacy in a more challenging F9 teratocarcinoma model remained transient^25^. EDA-CAR-T-treated tumors showed a delay in tumor growth by day 4 post-infusion, followed by the development of resistance as shown by exponential tumor growth 10 days after CAR T cell infusion (Fig. 1A-B). This initial response was characterized by marked EDA-CAR-T cell infiltration in the tumor and spleen, alongside a significantly higher frequency of intratumoral CD45⁺ immune cells and increased activation markers ICOS and CD137 in splenic CAR-T cells. However, by day 10, these response hallmarks were no longer observed suggesting a rapid loss of tumor control and unsustained CAR-T cell activity (Fig. S1, Table S2). To elucidate the mechanisms underlying limited therapeutic efficacy of EDA-CAR-T cells and the impact of the TME, we performed a scRNA-seq analysis on tumor samples collected at day 4 (peak of response) and day 10 (tumor progression) from animals treated with EDA-CAR-T cells, PSMA-CAR-T cells as control, together with CAR-T infusion products (IP) (Fig. 1C). Our integrated scRNA-seq dataset of the tumor samples comprised 84,171 high-quality cells across 15 independent biological replicates and included 24,485 CD45⁺ (*Ptprc^+^*) immune cells, 1,975 CAR-T cells (identified as *Ptprc^+^ Cd3e^+^ Gfp^+^*) and 57,711 (*Ptprc^-^*) non-immune/malignant cells. Unsupervised clustering delineated a highly heterogeneous TME, encompassing tumor cells, fibroblasts, and a diverse immune compartment including endogenous CD4⁺ and CD8⁺ T cells, NK cells and myeloid cells (Fig. 1D-F), that were annotated based on the expression of canonical markers (Fig. 1G, Fig. S2A-B, Table S3). This cellular heterogeneity was validated via flow cytometry (Fig. S2C). Quality control metrics demonstrated that samples contributed a balanced proportion of cells to each cluster across experiments and timepoints, confirming the absence of significant batch effects and ensuring a robust framework for downstream analysis (Fig. S3).

**Figure 1.**
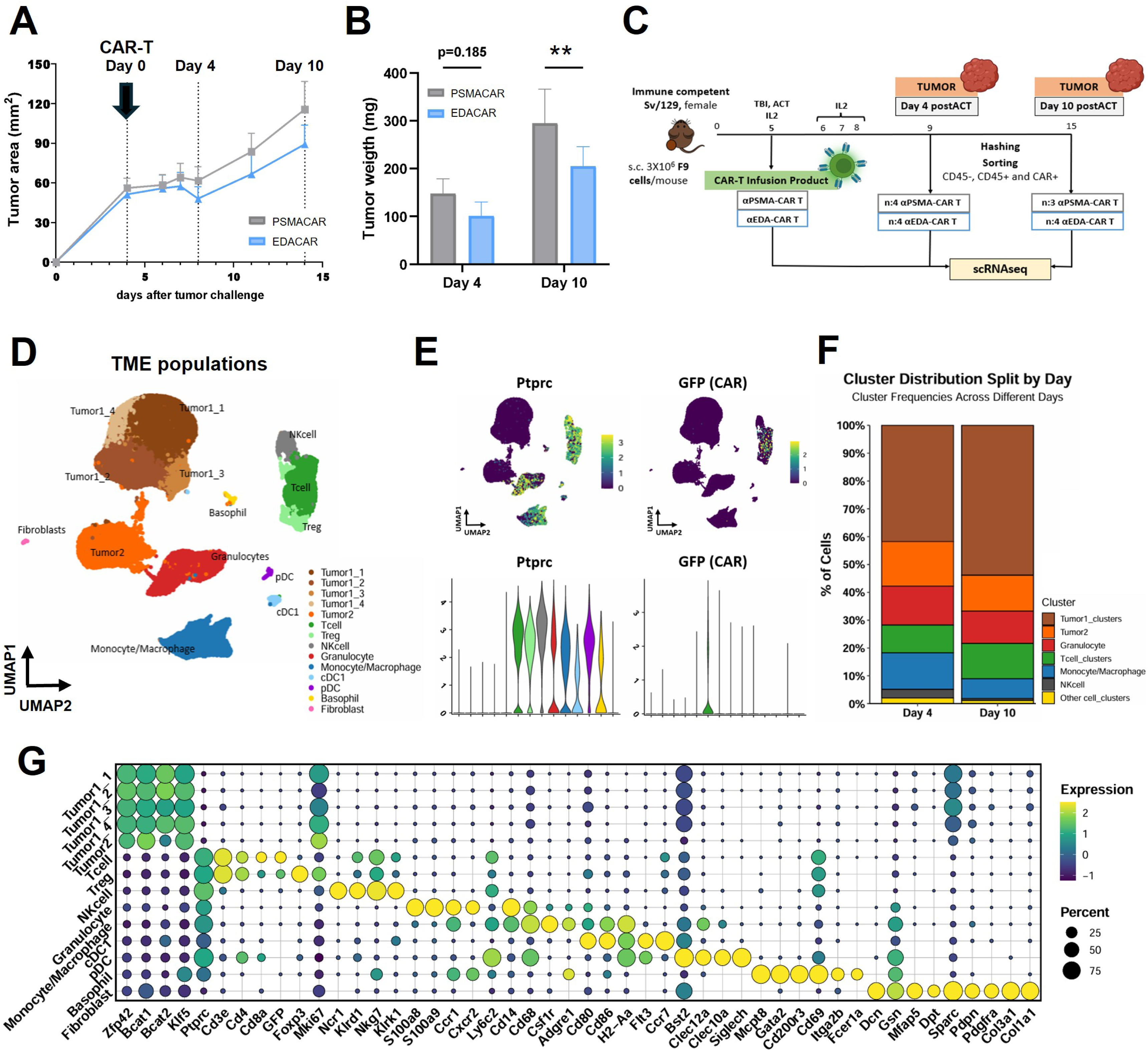
Comprehensive single-cell RNA-seq analysis of TME after treatment with EDA-CAR-T cells. **(A)** Tumor growth kinetics following treatment with EDA-CAR-T (n: 8) or control PSMA-CAR-T cells (n:7). Day 0 corresponds to the time of CAR-T infusion. **(B)** Tumor weights measured at day 4 (n:4 PSMA-CAR-T and EDA-CAR-T) and day 10 post-CAR-T infusion (n:3 PSMA-CAR-T and n:4 EDA-CAR-T). One-way ANOVA with Bonferroni’s multiple comparisons test. **(C)** Experimental design and workflow for scRNA-seq study (day 4, n:4 PSMA-CAR-T and EDA-CAR-T; day 10, n:3 PSMA-CAR-T and n:4 EDA-CAR-T; IP, n:1 PSMA-CAR-T and n:1 EDA-CAR-T). **(D)** UMAP visualization of major TME populations identified across all samples. **(E)** *Ptprc* (CD45) and GFP (CAR marker) relative expression across TME clusters shown on UMAP (top) and violin plots (bottom). **(F)** Relative abundance of major TME cell types at day 4 and day 10 post-CAR-T infusion. **(G)** Average expression of canonical marker genes used to define TME populations. Statistical significance is denoted as **P < 0.01. Bars representing the mean and SD are plotted (B and C). TBI, total body irradiation; ACT, adoptive cell transfer (CAR-T cell infusion); IL-2, recombinant human interleukin-2 support; s.c., subcutaneous tumor implantation; ANOVA, Analysis of Variance.

### Effector CD8^+^ EDA-CAR-T cell populations are enriched at peak of anti-tumoral response

To further characterize the phenotypic and functional dynamics of EDA-CAR-T cells following infusion, we performed sub-clustering analysis of endogenous T cells and CAR-T cells isolated from the TME dataset, integrated with IP samples (Fig. 2A). This analysis revealed distinct functional states, with CD8⁺ and CD4⁺ T cells organized along a developmental continuum encompassing memory populations, diverse effector subsets (co-expressing activation and exhaustion-associated markers), and transitional states with intermediate effector-memory profiles. In addition, we identified smaller clusters of highly proliferative T cells, regulatory T cells (Tregs), Th17-like CD4⁺ T cells, and cytotoxic γδ T cells (Fig. 2A-B, Fig. S4A-C). Notably, CAR-T cells were distributed across all identified T cell clusters, recapitulating the broad phenotypic heterogeneity of the endogenous T cell compartment within the TME (Fig. S4D-F).

**Fig 2.**
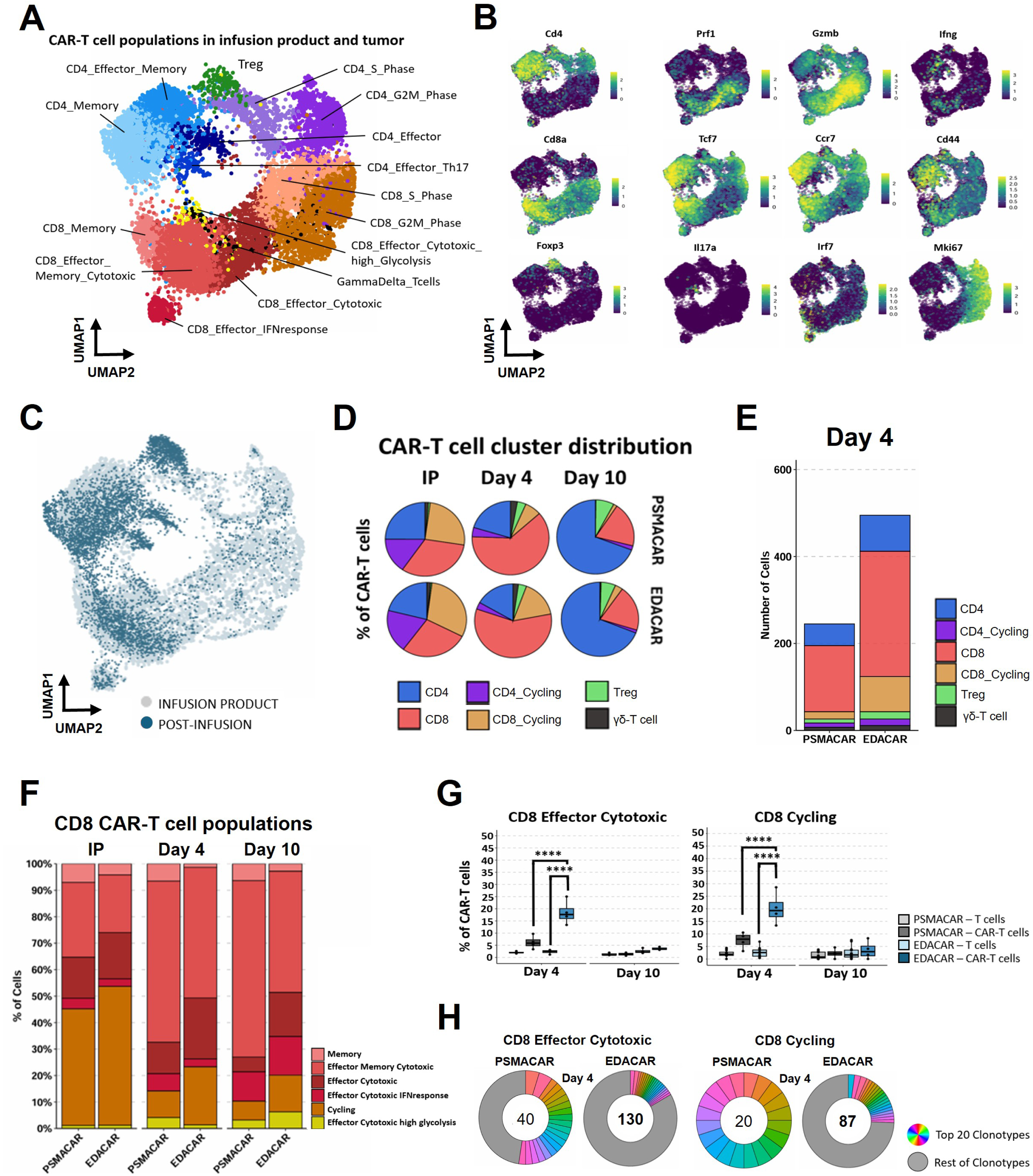
Effector CD8+ EDA-CAR-T cell populations are enriched at peak of anti-tumoral response. **(A)** UMAP visualization of CAR-T cell subclusters identified across the infusion product and post-infusion tumor samples. **(B)** Feature plots showing the expression of key identity and functional markers defining CAR-T cell subclusters. **(C)** UMAP projection showing CAR-T cells distribution across infusion product and post-infusion tumor samples. **(D)** Proportion of major CAR-T cell subsets (CD4⁺, CD8⁺, CD4⁺ cycling, CD8⁺ cycling, Treg, and γδ T cells) in infusion product (IP) and tumors at day 4 and day 10 post-infusion. **(E)** Absolute numbers of major CAR-T cell subsets isolated from tumors at day 4 after infusion. **(F)** Distribution of CD8⁺ CAR-T subclusters across the infusion product (IP), day 4, and day 10 post-infusion tumor samples. **(G)** Abundance of effector cytotoxic and cycling CD8⁺ CAR-T cell subsets at day 4 and day 10 post-infusion across biological replicates. Two-way ANOVA with Tukey’s multiple comparisons test. The box represents the interquartile range, the line indicates the median, and whiskers denote minimum to maximum values. **(H)** Clonotype diversity of effector cytotoxic and cycling CD8⁺ CAR-T cells at day 4 post-infusion. Statistical significance is denoted as ****Padj < 0.0001. ANOVA, Analysis of Variance.

We next examined the longitudinal evolution of EDA-CAR-T and PSMA-CAR-T cells following infusion. At the infusion stage, both products exhibited similar compositions, with enrichment in proliferative subsets (Fig. 2C-D). However, although tumor-infiltrating CAR-T cells from both conditions predominantly transitioned into CD8⁺ non-proliferative cells (Fig. 2D, Fig. S5A), EDA-CAR-T cells showed enhanced intratumoral infiltration at the peak of response (day 4) (Fig. 2E). Notably, a detailed characterization revealed that EDA-CAR-T cells displayed a pronounced skewing toward more differentiated effector states relative to PSMA-CAR-T cells (Fig. 2F). This shift was characterized by a significant enrichment of cytotoxic and proliferative CD8⁺ clusters, highlighting a distinct functional polarization of EDA-CAR-T cells within the TME (Fig. 2G). Moreover, this effector CAR-T cells enrichment was exclusive to the intratumoral EDA-CAR-T cells since it was not detected either in the IP nor within the endogenous T cell subsets (Fig. 2F, Fig. S5B). TCR sequencing further revealed a polyclonal expansion of effector populations, consistent with a broad, antigen-driven response rather than stochastic outgrowth of limited clones (Fig. 2H). However, at the stage of tumor progression (day 10), we observed that CD8⁺ CAR-T cell populations collapsed and both EDA-CAR-T and PSMA-CAR-T cell composition underwent a profound shift toward enrichment of CD4⁺ subsets, including Tregs (Fig. 2D). Interestingly, EDA-CAR-T cells at this stage were specifically enriched in polyclonal effector CD4⁺ cells (Fig. S5C-F). Importantly, this CD8-to-CD4 skewing was specific to the intratumoral CAR-T compartment, as IP and endogenous tumor-infiltrating T cells maintained a relatively stable CD4/CD8 ratio across both timepoints (Fig. S5G-H). Collectively, these findings delineate a distinctive tumor-specific and temporally dynamic evolution of EDA-CAR-T cell states within the TME that defines therapeutic outcome and highlight the central role of CD8⁺ effector EDA-CAR-T cell subsets in mediating the early antitumor response.

### EDA-CAR-T cells undergo transcriptional reprogramming towards dysfunction in the TME

To delineate the intrinsic molecular drivers of EDA-CAR-T cell functionality, we specifically analyzed transcriptional programs of effector cytotoxic CD8⁺ T cells (enriched population at peak of response) and effector CD4⁺ T cells (enriched population at disease progression). CD8⁺ EDA-CAR-T cells at day 4 showed significant enrichment of pathways associated with T cell activation, adhesion, degranulation, and cytotoxic function (Fig. 3A-B, Table S4&5). Moreover, pseudobulk GSVA confirmed the coordinated upregulation of effector programs across independent biological replicates (Fig. 3C, Table S7), which include key mediator genes such as *Ifng, Ccl5* or *Gzmk* (Fig. 3D). However, by day 10, these cells underwent a profound functional decline, downregulating activation, adhesion and proliferation pathways, concurrently with an enrichment terminal exhaustion program (Fig. 3A), characterized by the upregulation of exhaustion (*Lag3, Havcr2 and Pdcd1*) and apoptotic markers, alongside attrition of the effector machinery (Fig. 3C-D). Altogether, these findings indicate that while EDA-CAR-T cells initially achieve a potent cytotoxic state, they rapidly progress toward terminal exhaustion within the TME.

**Figure 3.**
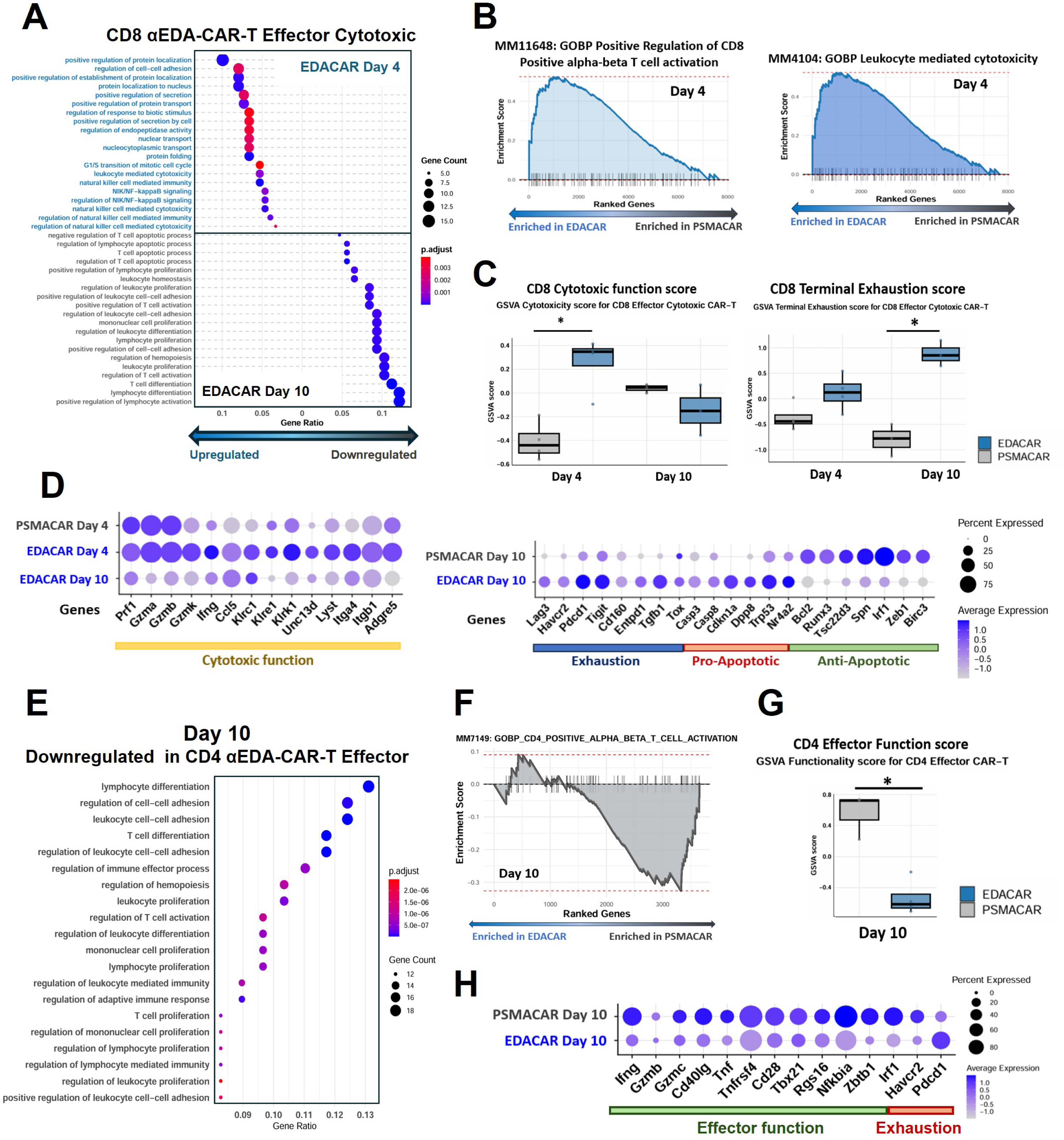
EDA-CAR-T cells suffer a widespread transcriptional reprogramming towards dysfunction in the TME. **(A)** Differential Gene Ontology enrichment analysis for biological functions of effector cytotoxic CD8⁺ EDA-CAR-T cells compared to control PSMA-CAR-T cells. Top: biological processes upregulated at day 4 post-infusion. Bottom: biological processes downregulated at day 10 post-infusion. **(B)** GSEA of curated gene signatures showing differential enrichment of CD8⁺ T cell activation (MM11648) and leukocyte-mediated cytotoxicity (MM4104) between effector cytotoxic CD8⁺ EDA-CAR-T and PSMA-CAR-T cells at day 4 post-infusion. **(C)** Pseudobulk GSVA scoring of effector cytotoxic CD8⁺ CAR-T cells at day 4 and day 10 using custom *cytotoxic function* (left) and *terminal exhaustion* (right) gene sets. **(D)** Average expression of key genes from the *cytotoxic function* (left) and *terminal exhaustion* (right) gene sets used for GSVA scoring in effector cytotoxic CD8⁺ CAR-T cells. **(E)** Differential Gene Ontology enrichment analysis for biological functions for down-enriched genes at day 10 post-infusion of effector CD4⁺ EDA-CAR-T cells compared to control PSMA-CAR-T cells. **(F)** GSEA of the curated CD4⁺ T cell activation signature (MM7149) comparing effector CD4⁺ EDA-CAR-T and PSMA-CAR-T cells at day 10. **(G)** Pseudobulk GSVA scoring of effector CD4⁺ CAR-T cells at day 10 using a custom *CD4 effector function* gene set. **(H)** Average expression of representative genes from the *CD4 effector function* gene set used for GSVA scoring in effector CD4⁺ CAR-T cells. Kruskal-Wallis test followed by Dunn’s post hoc test with Benjamini-Hochberg correction. Pairwise comparisons were performed using the Wilcoxon rank-sum test (C and G). The box represents the interquartile range, the line indicates the median, and whiskers denote minimum to maximum values (C and G). Statistical significance is denoted as *Padj < 0.05. GO, gene ontology; GOBP; gene ontology biological processes; GSVA, Gene Set Variation Analysis.

We next analyzed effector CD4⁺ EDA-CAR-T cells at day 10, which revealed a systemic collapse of functional programs. This subset showed negative enrichment of leukocyte-mediated immunity, differentiation, and adhesion pathways (Fig. 3E, Table S5), and significant deficit in helper and effector modules (Fig. 3F, Table S6). This dysfunctional state was characterized by the reduced expression of critical co-stimulatory and effector genes, including *Ifng, Cd40lg and Tbx21*, together with increased *Pdcd1* expression (Fig. 3G-H, Table S4). While analysis of the less differentiated effector-memory CD4⁺ EDA-CAR-T cell subset at day 4 showed early activation signatures (Fig. S6A-B, Table S5&6) and upregulation of effector genes, we observed that master Th1-lineage regulators (*Tbx21, Stat1*) were notably suppressed (Fig. S6C), accompanied by the precocious expression of the Th2-lineage driver *Gata3* (Fig. S6D), suggesting an uncomplete Th1-type activation state. These findings indicate that while CD4⁺ EDA-CAR-T cells undergo early priming, they may fail to consolidate a long-term functional differentiation program.

Crucially, the activation states observed at day 4 were not pre-imprinted during CAR-T cell manufacture. The transcriptional profiles of EDA- and PSMA-CAR-T cells in the IP were nearly indistinguishable, with no baseline enhancement of activation markers (Fig. S7A). In contrast, intratumoral EDA-CAR-T cells significantly upregulated effector programs relative to their IP in both CD8⁺ effector cytotoxic and CD4⁺ effector-memory subsets (Fig. S7A-C, Table S5&7), confirming that this functional reprogramming is a tumor-driven process. In summary, our results suggest that EDA-CAR-T cells undergo rapid transcriptomic reprogramming from early functional activation to widespread dysfunction within the TME, providing a mechanistic basis for the transient therapeutic efficacy.

### EDA-CAR-T cells remodel the TME but fail to induce an effective endogenous response

In addition to CAR-T intrinsic factors, we also evaluated extrinsic factors contributing to the transient efficacy of EDA-CAR-T therapy. Thus, we examined cell-to-cell interactions between CAR-T cells and endogenous TME (Fig. S8, Table S8). Cell-cell communication analysis revealed a robust IFNγ-mediated signaling network at day 4 (Fig. 4A), with unique interactions between CD8⁺ EDA-CAR-T cells with macrophages and fibroblasts that were absent in the control group (Fig. 4B), indicative of an early EDA-CAR-T-specific inflammatory response. Among macrophages, the endothelial-migratory subset likely represents a primary responder to EDA-CAR-T-derived IFNγ due to its spatial proximity to the EDA-enriched stromal-vascular niche^26^. These macrophages from EDA-CAR-T-treated tumors were significantly enriched at day 4 in pathways for T-cell engagement, including MHC-I, CD48, ICAM, and PECAM, with several predicted interactions being EDA-CAR-T-specific (CD80, PECAM, and ITGB2) (Fig. 4C-E). However, CD80/CD86-centered networks integrated both CD28 and CTLA-4 from EDA-CAR-T cells, suggesting potential simultaneous co-stimulatory and co-inhibitory signaling, which further extends to additional antigen-presenting cells (APC) like cDC1 cells, monocytes and other macrophage subsets (Fig. 4E-F, Fig. S4F). Beyond macrophages, we observed that cDC1 cells in EDA-CAR-T-treated tumors also established strengthened communication with CAR-T cells at day 4, driven by CD40LG-CD40 interactions (CD4⁺ CAR-T) and FLT3L signaling (CD8⁺ CAR-T) (Fig. S9A-B). This was accompanied by upregulation of antigen-processing genes (*H2-Aa, Tap1, Psmb2*), immunostimulatory molecules (*Icosl, MHC-I, MHC-II*), and immunoregulatory genes including *Pdcd1lg2* and *Lgals1* (Fig. S9C-F, Table S4). Again, although ICOS-ICOSL interactions were specific to EDA-CAR-T cells, it was predicted to interface with both CD28 and CTLA-4 (Fig. S9G-H). Together, these results suggest that EDA-CAR-T cells successfully engage the myeloid compartment, but the resulting communication network integrates co-stimulatory and co-inhibitory signals that may simultaneously promote and restrain T cell activation.

**Figure 4.**
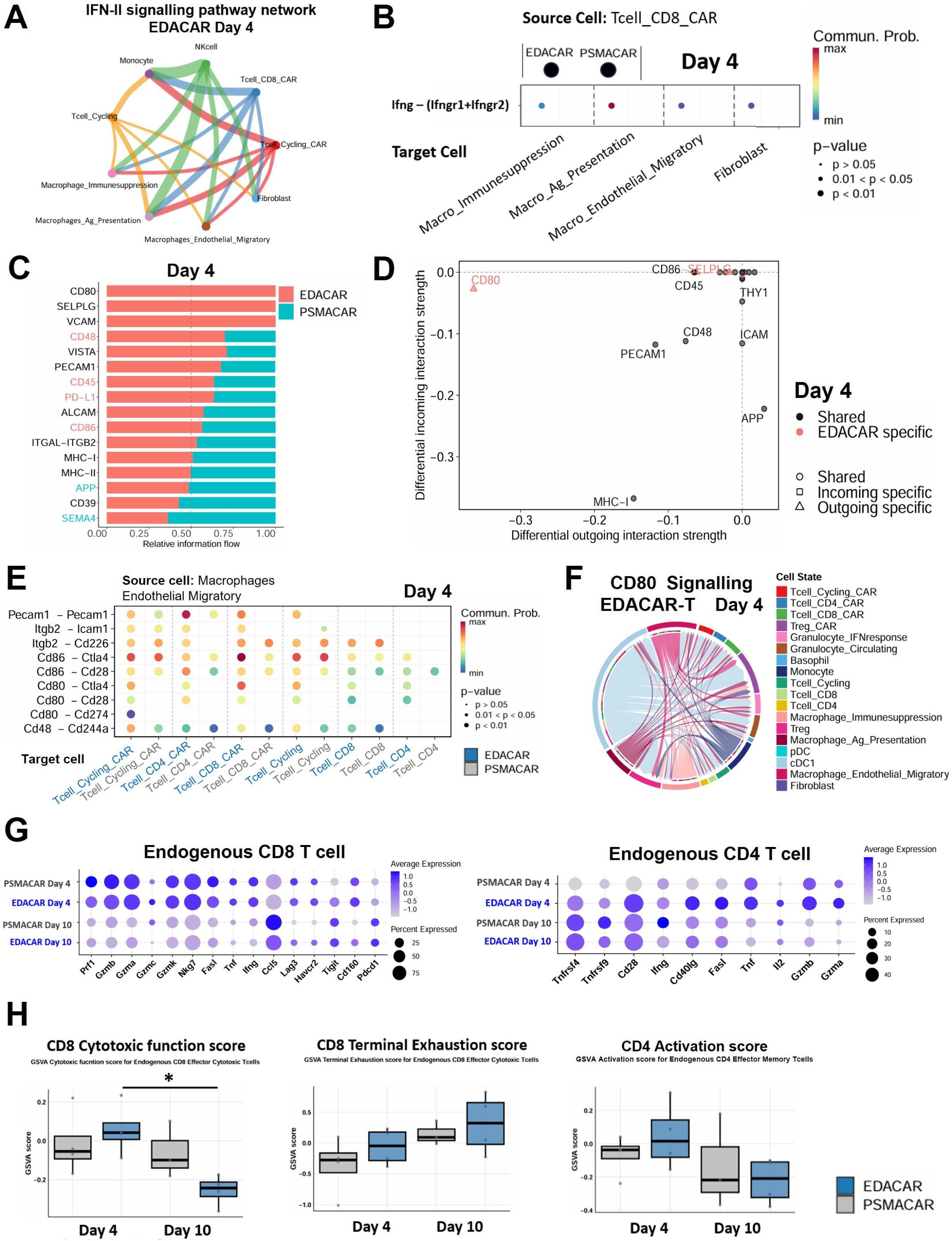
EDA-CAR-T cells fail to promote an effective Th1-like immune response. **(A)** Circle plot depicting IFN-II (IFNγ, Ifng) signaling networks in EDA-CAR-T-treated tumors at day 4 post-infusion. **(B)** Differential IFNγ ligand-receptor interactions from CD8⁺ EDA-CAR-T cells (source) compared to control PSMA-CAR-T cells at day 4 post-infusion. For each target cell population, the left dot represents the interaction probability in EDA-CAR-T-treated tumors, whereas the right dot represents the interaction probability in PSMA-CAR-T-treated control tumors. **(C)** Differential overall information flow within the inferred *cell-cell contact* signaling network for Macrophages_Endothelial_Migratory (source) from EDA-CAR-T-treated tumors compared to control PSMA-CAR-T-treated at day 4 post-infusion. **(D)** Differential incoming and outgoing signaling strength within the *cell-cell contact* network for Macrophages_Endothelial_Migratory (source) from EDA-CAR-T-treated tumors compared to control PSMA-CAR-T-treated at day 4 post-infusion. **(E)** Differential PECAM1, ICAM, CD80, CD86 and CD48 ligand-receptor interactions between Macrophages_Endothelial_Migratory (source) and T cells and CAR-T cells (target) from EDA-CAR-T-treated tumors compared to control PSMA-CAR-T-treated at day 4 post-infusion. **(F)** Chord plot showing CD80-mediated signaling networks across the TME in EDA-CAR-T-treated tumors at day 4 post-infusion. **(G)** Average expression of key activation, effector and exhaustion genes in endogenous CD8⁺ and CD4⁺ T cells across treatment groups. **(H)** Pseudobulk GSVA scoring of endogenous T cells at day 4 and day 10 post-infusion: effector cytotoxic CD8⁺ T cells scored using *cytotoxic function* (left) and *terminal exhaustion* (middle) gene sets, and effector-memory CD4⁺ T cells scored using a custom *CD4 activation* gene set (right). Kruskal-Wallis test followed by Dunn’s post hoc test with Benjamini-Hochberg correction. Pairwise comparisons were performed using the Wilcoxon rank-sum test. The box represents the interquartile range, the line indicates the median, and whiskers denote minimum to maximum values. Statistical significance is denoted as *Padj < 0.05.

To determine if this early remodeling translated into a broader anti-tumor response, we assessed the functional state of endogenous T cells. Strikingly, CD8^+^ T cells from EDA-CAR-T-treated animals showed no significant functional activation compared to controls. They failed to upregulate cytotoxic programs or *Ifng* expression, instead maintaining high expression of exhaustion markers such as *Lag3* and *Havcr2*. While effector-memory CD4⁺ T cells showed a transient upregulation of activation and effector genes (*Cd40lg, Fasl, Il2*) at day 4, this response rapidly decayed by day 10 (Fig. 4G). Moreover, the robust effector modules defining the EDA-CAR-T cells at day 4 were almost entirely absent in the endogenous T-cell compartment (Fig. 4H, Table S6). Collectively, these findings demonstrate that while EDA-CAR-T cells can successfully remodel the TME by engaging the myeloid compartment, the resulting bystander T cell activation is incomplete and lacks sustained functional reprogramming.

### EDA-CAR-T cells exhibit increased sensitivity to inhibitory signals from the TME

To identify TME features driving EDA-CAR-T cell loss and suppression of endogenous immune response, we profiled immunosuppressive hallmarks across all major TME cell compartments. First, differential gene expression analyses identified tumor cells as a major source of metabolic pressure, with high expression of glycolytic enzymes and nutrient transporters (*Ldha, Slc2a1, Slc1a5*), as well as angiogenic factors (*Vegfa, Vegfb*), consistent with a central role in metabolic competition and angiogenesis. Hypoxia-associated gene expression (*Hif1a*) was broadly elevated across the TME, suggesting a hypoxic microenvironment. Furthermore, we observed upregulation of extracellular matrix components (*Col1a1, Fn1, Mmp9*), TGF-β signaling (*Tgfb1*), and immune-evasion pathways, including reduced MHC-I (*H2-K1*), increased non-classical MHC-I (H2-T23), checkpoint ligands (*Cd274, Pdcd1lg2*), metabolic suppressors (*Ptgs2, Arg1, Entpd1*), immunosuppressive chemokines (*Ccl8, Ccl22, Cxcl12*), and galectins (*Lgals1, Lgals3, Lgals9*) (Fig. S10A-B). Collectively, these data delineate a hostile TME governed by overlapping physical, metabolic, and biochemical barriers that likely impair CAR-T cell persistence and limit effective anti-tumor immunity (Fig. S10C).

We next investigated the interactome directly targeting CAR-T compartment. Global communication profiling revealed that CD8⁺ EDA-CAR-T cells at day 4 engaged in a significantly higher number of interactions compared to PSMA-CAR-T cells (Fig. 5A). At day 4, EDA-CAR-T cells showed enrichment for PD-L1, PD-L2 and TGFβ signaling pathways (Fig. S11A). PD-L1 signals from granulocytes, endothelial-migratory macrophages, and cDC1s converged on CD8⁺ and cycling CAR-T cells, alongside an autocrine PD-L2 loop within CAR-T subsets (Fig. S11B-C). Strikingly, although macrophages, cDC1 and pDCs were major sources of TGFβ signaling within the TME, EDA-CAR-T cells also contributed (Fig. S11D). TGFβ signals were strongly directed toward macrophages and cDC1, raising the possibility that CAR-T-derived TGFβ interactions may suppress APC subsets required to sustain a productive anti-tumoral response (Fig. S11E-F). Consistently, enhanced TGFβ interactions with endothelial-migratory macrophages persisted at days 4 and 10 (Fig. S11G), suggesting a negative feedback loop limiting APC immunostimulatory potential. Together, these results reveal a multilayered suppressive architecture. The convergence of extrinsic checkpoint signaling and intrinsic autocrine feedback (via TGFβ and PD-L1/PD-L2) would provide a mechanistic basis for the transient efficacy and eventual functional decline of EDA-CAR-T cell therapy.

**Figure 5.**
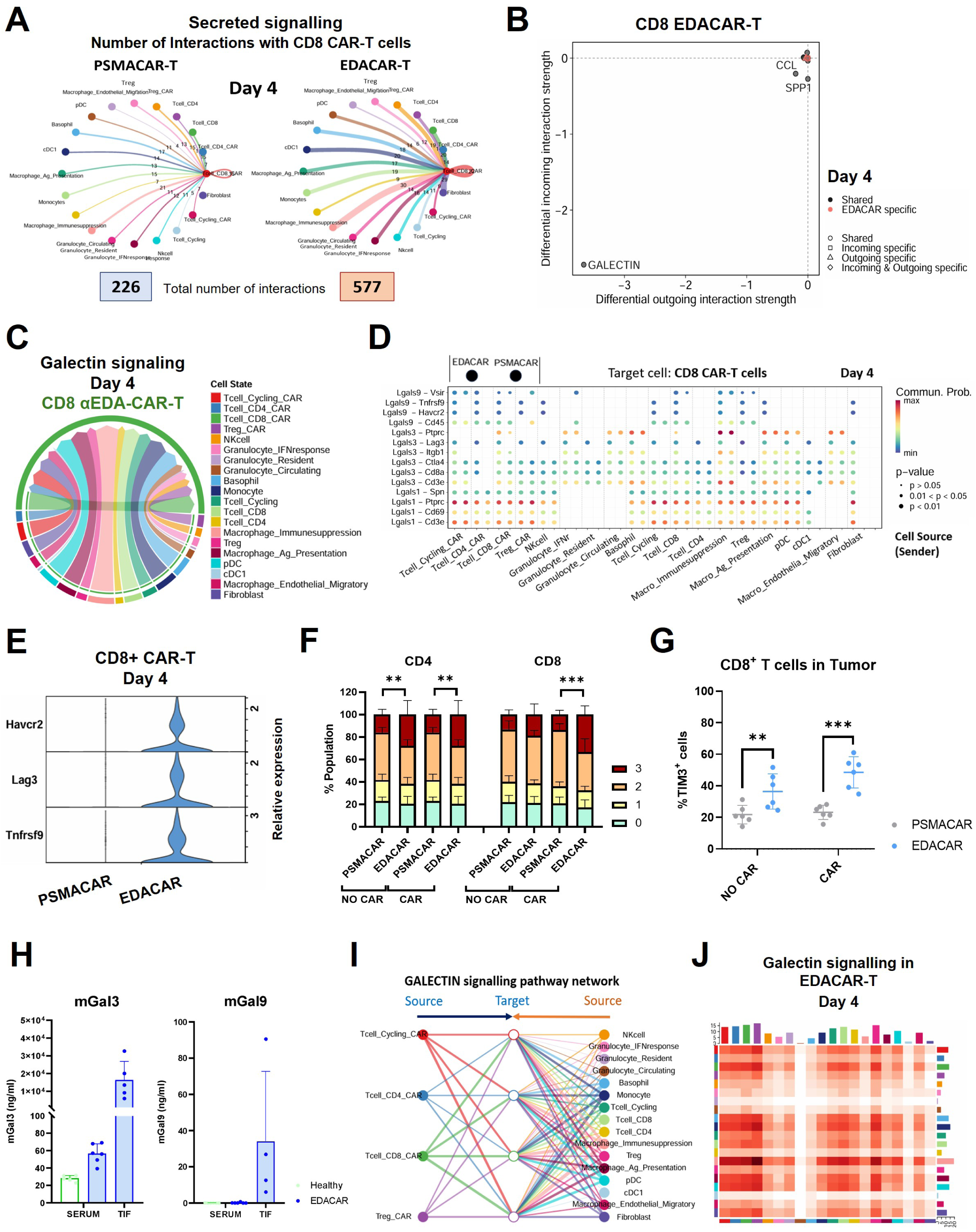
EDA-CAR-T cells exhibit increased sensitivity to galectin-mediated inhibitory signals. **(A)** Circle plot showing the total number of inferred *Secreted Signaling* interactions received by CD8⁺ CAR-T cells in PSMA-CAR-T- and EDA-CAR-T-treated tumors at day 4 post-infusion. **(B)** Differential incoming and outgoing signaling strength within the *Secreted Signaling* network for CD8⁺ EDA-CAR-T cells (target) compared to control PSMA-CAR-T cells at day 4 post-infusion. **(C)** Chord diagram illustrating galectin-mediated interactions targeting CD8⁺ EDA-CAR-T cells across the TME at day 4 post-infusion. **(D)** Differential ligand--receptor interactions for the Galectin pathway between CD8⁺ CAR-T cells (target) and the TME populations (source), showing EDA-CAR-T-specific inhibitory axes relative to PSMA-CAR-T controls. For each cell source population, the left dot represents the interaction probability in EDA-CAR-T-treated tumors, whereas the right dot represents the interaction probability in PSMA-CAR-T-treated control tumors. **(E)** Relative expression of the galectin-associated inhibitory receptors *Havcr2* (TIM3), *Lag3*, and *Tnfrsf9* in CD8⁺ CAR-T cells at day 4 post-infusion. **(F)** Frequency of exhaustion marker co-expression in endogenous T cells (NO CAR) and CAR-T cells (CAR) from tumors of PSMA-CAR-T- or EDA-CAR-T-treated mice at day 4 post-infusion, analyzed by flow cytometry (n = 6 biological replicates per group). Exhaustion markers assessed were LAG-3, TIM-3, and PD-1. Stacked bars represent the proportion of cells expressing 0, 1, 2, or 3 exhaustion markers, corresponding to the cumulative number of markers co-expressed per cell. **(G)** Percentage of TIM3 positive T cells (NO CAR) and CAR-T cells (CAR) in the tumor from PSMA-CAR-T- or EDA-CAR-T-treated mice at day 4 post-infusion analyzed by flow cytometry. **(H)** Concentration of mouse soluble galentin-3 (mGal3) and -9 (mGal9) in serum (Healthy n:6; EDA-CAR n:6 biological replicates for mGal3 and mGal9) and tumor interstitial fluids (EDA-CAR n:5 biological replicates for mGal3 and n:4 biological replicates for mGal9). **(I)** Hierarchical plot showing galectin signaling architecture in EDA-CAR-T-treated tumors at day 4. Filled nodes represent signaling sources; open nodes represent targets. Edge colors correspond to the source cell type. **(J)** Distribution and strength of galectin-mediated interactions across all TME populations at day 4 post-infusion; color-coding of cell types matches the legend used in panel (C). Two-way ANOVA with Tukey’s multiple comparisons test (F and G). Bars and lines representing the mean and SD are plotted (F-H). Statistical significance is denoted as **P < 0.01 and ***P < 0.001. ANOVA, analysis of variance; mGal3 and 9, mouse soluble galectin-3 and -9.

To broaden our assessment, we interrogated additional secreted-inhibitory signals. Among these, galectin pathway emerged as the most differentially enriched signaling network (Fig. 5B-C). Ligand-receptor analysis identified EDA-CAR-T-specific inhibitory axes mediated by galectin-1, -3 and -9 at day 4, including Galectin-3/LAG3, Galectin-9/HAVCR2 and Galectin-9/TNFRSF9 (Fig. 5D). Notably, these interactions were absent in PSMA-CAR-T cells and correlated with the transcriptional upregulation of *Lag3, Havcr2*, and *Tnfrsf9* in EDA-CAR-T cells (Fig. 5D-E). Flow cytometry confirmed an increased exhaustion profile of EDA-CAR-T cells, with specific upregulation of TIM-3 (Fig. 5F-G), while soluble galectin-3 and -9 were detected in tumor interstitial fluids (Fig. 5H). These data suggest that antigen-driven activation renders EDA-CAR-T cells uniquely sensitized to galectin-mediated inhibition. Endothelial-migratory macrophages and cDC1s emerged as key nodes in this suppressive network, reinforcing their dual role in providing both activation and inhibitory cues (Fig. 5D). Soluble galectin signaling extended beyond CD8⁺ EDA-CAR-T cells to CD4⁺ and cycling EDA-CAR-T subsets and endogenous T cells, broadening suppressive pressure across the TME (Fig. 5I-J). Taken together, these results identify galectin-mediating signaling networks as major inhibitory axis potentially driving the EDA-CAR-T cells decline within the TME.

## DISCUSSION

Unlike blood cancers, solid tumors impose critical barriers, including restricted tissue infiltration, chronic antigen exposure and immunosuppressive TME, that impairs CAR-T cell function and persistence. Consequently, the mechanisms underlying CAR-T efficacy in hematological malignancies cannot be directly translated to solid tumors, highlighting the need for detailed mechanistic studies in relevant preclinical and clinical settings. We leveraged single-cell RNA-seq to comprehensively characterize the cellular diversity, functional states and intercellular communication networks of CAR-T cells within the TME, providing a high-resolution framework to identify molecular mechanisms driving CAR-T cell functionality in solid tumors.

Our single-cell RNA-seq analysis reveals that peak antitumoral efficacy correlates with improved intratumoral EDA-CAR-T cell infiltration and enrichment of polyclonal CD8⁺ effector and effector-memory EDA-CAR-T cells. These CAR-T cells display robust activation and cytotoxic transcriptional programs, characterized by high expression of effector genes (*Ifng*, *Cd44* and *Cxcr6*) alongside downregulation of memory-associated genes including (*Tcf7, Ccr7* and *Sell*). This profile defines a highly activated, tumor-engaged effector T cell state and aligns with clinical and preclinical evidence linking disease control in solid tumors to the intratumoral accumulation of cytotoxic CD8⁺ CAR-T cell subsets^26,27^. Moreover, our data indicates that as therapeutic efficacy declines, this CD8⁺ CAR-T population rapidly progresses towards terminal exhaustion programs and contracts. This marked loss of tumor-infiltrating effector CD8⁺ EDA-CAR-T cells strongly suggests that poor persistence could represent a principal mechanistic determinant of therapeutic failure. Limited CAR-T cell persistence is widely recognized as a major barrier to therapeutic efficacy in solid tumors. Rising number of studies have demonstrated that enhancing CAR-T cell persistence can directly translate into more sustained antitumor responses^28–34^. Notably, these studies highlight that reinforcing activation and memory formation in lymphoid tissues, while simultaneously promoting effector infiltration and functionality within tumors, is key for achieving durable CAR-T cell responses, paving the way for the development of improved CAR-T cell strategies in solid tumors. Our results reveal that EDA-CAR-T and endogenous T cells residing in the spleen adopt a memory state but show low long-term activation phenotype, suggesting poor priming or reinforcement in secondary lymphoid organs. This insufficient lymphoid activation may therefore contribute to limited long-term persistence and loss of therapeutic efficacy observed with EDA-CAR-T therapy.

In addition to limited persistence, our data indicates that CD8⁺ EDA-CAR-T cells upregulate multiple exhaustion-associated markers, including *Pdcd1, Havcr2, Tigit, Lag3, Ctla4, Gzmk* and *Tnfrsf9*. This transcriptional profile closely mirrors observations from both clinical and preclinical studies of solid tumors, where intratumoral CD8⁺ CAR-T cells and endogenous TILs are consistently enriched in exhausted subsets^26,35^. These findings suggest that exhaustion represents a prevalent and defining feature of tumor-infiltrating CD8⁺ T cell states. Importantly, this exhaustion signature has been widely associated with CAR-T cell functional impairment and poor prognosis in patients^36,37^. Interestingly, Sawada et al. showed that PD-1+ Tim3+ tumor-infiltrating CD8 T cells sustain the potential for IFN-γ production but lose cytotoxic activity in ovarian cancer, indicating that these exhausted CAR-T cells could still retain partial effector capacity^36^. Consistent with this idea, our results show that terminally exhausted cytotoxic CD8⁺ EDA-CAR-T cells maintain high Ifng expression at day 10 post-infusion, suggesting preserved effector function despite acquisition of a terminal exhaustion program. Strikingly, recent studies have demonstrated that such exhausted CD8⁺ T cell states can be functionally rescued. For example, administration of Fc-IL-4 and Fc-IL-10 has been shown to directly act on PD-1⁺TIM-3⁺ terminally exhausted CD8⁺ T cells, enhancing their cytotoxic capacity and promoting accumulation of functional CD8⁺ populations within tumors, thereby improving the efficacy of adoptive T cell transfer and immune checkpoint blockade therapies^38,39^. Collectively, these observations support that the exhausted-like phenotype observed in intratumoral CD8⁺ CAR-T cells may reflect an adaptive response to chronic antigen stimulation or immunosuppression within the TME rather than an irreversible loss of function. From this perspective, exhaustion programs could represent a dynamic and potentially reversible state that may be therapeutically exploited to restore CAR-T cell functionality in solid tumors.

The cancer-immunity cycle is widely recognized as a central framework underlying the development of effective antitumor immune responses^40^. Accumulating clinical and preclinical evidence indicates that anti-tumor efficacy of CAR-T cell therapies in solid tumors is accompanied with the polyclonal expansion, tumor infiltration, and activation of endogenous CD4⁺ and CD8⁺ T cells, underscoring the importance of engaging host immunity beyond the transferred CAR-T compartment^26,27^. Our data show that EDA-CAR-T cells could ignite early endogenous immune activation within the TME. Specifically, EDA-CAR-T cell therapy could engage endothelial-associated migratory macrophages and cDC1 cells, enhancing their antigen-presenting capacity and promoting immune crosstalk through upregulation of co-stimulatory molecules (MHC-I, MHC-II, CD80, CD86, ICOS) and adhesion-related pathways (PECAM1, ITGB2). Many studies have demonstrated that reinforcing myeloid and innate immune activation can markedly enhance CAR-T cell therapeutic efficacy in solid tumors. These strategies promote robust Th1 polarization, increased NK cell and T cell infiltration and activation, maturation of dendritic cells, and skewing of macrophages toward M1-like phenotypes, while concomitantly reducing immunosuppressive populations such as M2-like macrophages, regulatory T cells, and tolerogenic DC cells^27,41–48^. Collectively, these observations highlight that, in solid tumors, CAR-T cell efficacy may critically depend not only on direct tumor cytotoxicity but also on the capacity to reshape the TME and engage endogenous immunity. Our results reveal that EDA-CAR-T cell therapy induces early remodelling of antigen-presenting cell networks, which is subsequently associated with enhanced infiltration of endogenous CD8⁺ T cells and activation of CD4⁺ T cells, as evidenced by increased expression of effector molecules including *Ifng, Cd40lg, Fasl, Il2,* and *Tnf*. However, despite this initial immune stimulation, these responses did not translate into functional activation of endogenous CD8⁺ T cells and were accompanied by a long-term increased accumulation of regulatory T cell subsets in the TME. Together, these data indicate that EDA-CAR-T therapy fails to effectively propagate a sustained effector Th1-type antitumoral immune response. The inability to amplify and sustain a coordinated Th1-driven immune program may represent a key limitation of EDA-CAR-T therapy, offering a mechanistic explanation for its poor long-term antitumoral efficacy and supporting the rationale for additional immune-stimulatory approaches.

Our transcriptomic profiling across TME cell populations revealed hallmarks of hypoxia, metabolic competition, extracellular matrix remodelling and immunosuppression. These mechanisms have been extensively characterized in the literature as impairing CAR-T cell infiltration, function, and persistence in solid tumors, collectively defining the solid TME as a hostile and immunosuppressive milieu shaped by multiple, overlapping inhibitory mechanisms^11,49^. Accordingly, an increasing body of work has demonstrated that CAR-T cells can be rationally engineered to overcome such hostile TME conditions, resulting in improved antitumoral performance in solid tumors^50–61^. Together, it reinforces the idea that CAR-T failure in solid tumors is not driven by a single suppressive axis, but rather by the cumulative impact of a complex and redundant inhibitory landscape. Building on these observations, our data suggests that EDA-CAR-T cell activation may further exacerbate immunosuppressive pressure within the TME through engagement of soluble galectin-mediated axes TIM3-GAL9, TNFSFR9-GAL9 and LAG3-GAL3. These pathways have been proved to limit T cell and CAR-T cell functionality by promoting apoptosis of effector Ifng-secreting subsets^62–67^. While an increasing body of evidence correlates elevated levels of secreted galectins to poor prognosis in solid tumors, such as galectin-9 in pancreatic ductal adenocarcinoma^68,69^, several groups have shown that targeting TIM3-GAL9 axis in CAR-T cells improves anti-tumor efficacy and persistence^70,71^. Collectively, these observations support a central role for secreted galectins in reinforcing immune suppression and driving CAR-T and T cell dysfunction in solid tumors and highlight that effective CAR-T therapies for solid tumors will likely require not only enhanced tumor infiltration and activation, but also intrinsic resistance to dominant immunosuppressive cues within the TME.

In the present study, we have shown that scRNA-seq provides a powerful tool to decode the cellular diversity and functional dynamics of CAR-T cells within the complex TME. Our analysis highlights that understanding CAR-T cell function at single-cell resolution in the context of the TME is key to dissect integrated molecular mechanisms of therapeutic efficacy and decline. Beyond elucidating molecular pathways that govern CAR-T cell performance, this approach enables the identification of novel target candidates for modulation, thereby paving the way for the design of improved CAR-T therapies for solid tumors.

## DECLARATIONS

### Ethics approval

All animal handling and tumor experiments were approved and conducted under the institutional guidelines of our institutional ethics committee (Ref: 031-24) and following the European Directive 2010/63/EU.

### Authors’ contributions

RRF conceived the study, designed and performed *in vivo* experiments, conducted bioinformatic analyses, interpreted the data, and wrote the manuscript. PJL contributed to experimental design and data discussion. MECC provided bioinformatic analysis support. LJU, MGS and SCP contributed to data interpretation and results discussion. PSMU and PA performed scRNA-seq experiments. DA and AL conducted flow cytometry sorting and provided technical expertise and advisory support. SRD and RMT provided technical and experimental support. MG performed viral productions and provided *in vivo* experimental support. MH contributed to results discussion. JJL, FP, TL, and JRRM conceived and supervised the study, contributed to experimental design and data interpretation, wrote the manuscript and secured funding. All authors reviewed, edited, and approved the final manuscript.

## Acknowledgements and funding

This work was supported by the Government of Navarra Department of Health (GN2023/08 and GN2024/04). Spanish Ministry of Science, Innovation and Universities, grants PID2021-128283OA-I00, PID2022-137914OB-I00, PID2024-156683OB-I00 and PDC2025-166182-I00, funded by MICIU/AEI/10.13039/501100011033 and by FEDER, UE. Instituto de Salud Carlos III (ISCIII) TRANSCAN2022-784-114 (AC22_1/00006); Red de Terapias Avanzadas RICORS TERAV Plus (RD24/0014/0010), and Centro de Investigación Biomédica en Red de Cáncer (CIBERONC; CB16/12/00489, CB16/12/00369). The European Commission (EASYGEN, HORIZON-JU-IHI-2024-07, grant agreement No 101194710). Scientific Foundation of the Spanish Association Against Cancer (FC AECC) (PRYGN259200HERN). “La Caixa” Foundation under the project code LCF/PR/HR24/52440011. Paula and Rodger Riney Foundation. Alberto Palatchi Foundation.

RRF was supported by FPU fellowship (FPU2023/00653). PJL was supported by AECC postdoctoral fellowship (POSTD258767JUST) funded by the Scientific Foundation of the Spanish Association Against Cancer (FC AECC). PAR was supported by Ayudas para contratos de Personal Técnico de Apoyo (PTA) 2021 (PTA2021-020262-I). LJU acknowledges support from a WIT grant (Marie Skłodowska-Curie Actions, Horizon 2020). SCP was supported by FPU fellowship (FPU2023/00439) and from a CIMA AC fellowship. MH was supported by grant RYC2021-033127-I funded by MICIU/AEI/10.13039/501100011033 and by FEDER, UE.

The funders had no role in the design of the study; the collection, analysis, or interpretation of data; the writing of the manuscript; or the decision to submit the work for publication.

## Patient consent for publication

Not applicable

## Availability of data and material

All data needed to evaluate the conclusions in the paper are present in the paper and/or the Supplementary Materials. The scRNA-seq data generated in this study have been deposited in the GEO database (GSE325499).

