## Supplemental Material for "Rapid and dynamic reprogramming within the tumor microenvironment drives EDA-CAR-T dysfunction and compromised therapeutic efficacy in solid tumors"

### SUPPLEMENTARY MATHERIALS AND METHODS

**Mice**

Female 129Sv mice (10-12 weeks old) were obtained from Janvier Labs (Le Genest-Saint-Isle, France) and housed under specific pathogen-free conditions at the animal facility of the Centro de Investigación Médica Aplicada (CIMA, Pamplona). All animal procedures were approved by the local ethical committee and performed in accordance with institutional guidelines.

**Cell culture**

The murine teratocarcinoma F9 cell line (ATCC) was cultured in 0.1% Porcine Gelatin-coated flasks (Sigma) with complete RPMI 1640 medium supplemented with GlutaMAX™ (1x), 10% fetal bovine serum, 1× penicillin-streptomycin, 2 mM L-glutamine, 10 mM HEPES buffer, 1× MEM non-essential amino acids, 1 mM sodium pyruvate, 50 µM β-mercaptoethanol and 2.5 µg/mL amphotericin B (fungizone). The Platinum Ecotropic (Plat-E) (ATCC) cell line was cultured in complete DMEM 1640 medium supplemented with GlutaMAX™ (1x), 10% fetal bovine serum, 1× penicillin-streptomycin, 2 mM L-glutamine, 10 mM HEPES buffer, 1× MEM non-essential amino acids, 1 mM sodium pyruvate, 50 µM β-mercaptoethanol and 2.5 µg/mL amphotericin B (fungizone); and the selection antibiotics puromycin (100 µg/mL) and blasticidin (10 µg/mL). All cell lines were cultured at 37 °C in a humidified atmosphere containing 6.5% $CO_2$.

**Mouse T cell isolation**

Murine $CD4^+$ and $CD8^+$ T cells were obtained from the spleens and lymph nodes of female Sv129 mice. To obtain single-cell suspensions of T cells, organs were first homogenized through a cell strainer in ACK lysis buffer and passed through a magnetic column labelled with negative selection microbeads following manufacturers' instructions. $CD4^+$ and $CD8a^+$ T cell isolation kits (Miltenyi Biotec) were used to isolate mouse T cells. Purified lymphocytes were cultured in complete RPMI 1640 medium supplemented with GlutaMAX™ (1x), 10% FBS, 1x penicillin-streptomycin, 2 mM L-glutamine, 10 mM HEPES buffer, 1xMEM non-essential amino acids, 1 mM sodium pyruvate, 50 µM β-mercaptoethanol and 2.5 µg/mL amphotericin B (fungizone); and 100 IU/mL recombinant human interleukin-2 (rhIL-2).

**Viral vectors and virus production**

The chimeric EDA-CAR used to generate the murine CAR-T cells is composed of the anti-EDA F8 scFv and a murine 4-1BB-CD3ζ expression cassette linked through a F2A self-cleaving peptide sequence to eGFP. The PSMA-CAR, used as an irrelevant CAR, included the anti-human PSMA scFv obtained from mouse hybridoma J591, that was cloned in the same expression cassette. All constructs were synthesized by GenScript. For retroviral production, $7x10^5$ Plat-E cells were seeded per well in 6-well plates and transfected 24 h later with 5 µg of retroviral plasmid DNA along with 2.5 µg pCL-Eco plasmid DNA using lipofectamine 2000 for 6 hours in antibiotic-free medium. Retroviral supernatants were collected at 48 and 72 hours after transfection and filtered through 0.45-µm filters.

**Lymphocyte transduction**

Purified murine $CD4^+$ and $CD8^+$ T cells were activated using anti-CD3/CD28 Dynabeads at a bead-to-cell ratio of 1:2 for 24 h and at a cell density of $1x10^6$ cells/mL in complete RPMI supplemented with 100 IU/mL rhIL-2. T cells were transduced with retroviral supernatants in the presence of 10 µg/mL protamine sulfate and at a cell density of $0.25x10^6$ cells/mL by spinoculation at 1,300 × g for 90 min at 32 °C. Transduction was repeated 24 h later. T cells were expanded in complete RPMI supplemented with 100 IU/mL rhIL-2 until day 5 and used for downstream assays. Transduction efficiency was assessed 3 days after second retroviral transduction by flow cytometry based on reporter gene expression.

**In vivo tumor model and adoptive cell transfer**

Female 129Sv mice (10-12 weeks old) were subcutaneously injected in the right flank with $3x10^6$ F9 murine teratocarcinoma cells. Five days after tumor implantation, mice received lymphodepleting total-body γ-irradiation (4 Gy; exposure time: 42 s). Immediately after irradiation, mice were intravenously infused with $1x10^7$ CAR-T cells ($GFP^+$), consisting of a 1:1 mixture of $CD4^+$ and $CD8^+$ T cells. Recombinant human IL-2 (20,000 IU per mouse) was administered intraperitoneally once daily for four consecutive days following T cell infusion.

**Tumor and spleen collection and single-cell suspension preparation**

At days 4 and 10 post-adoptive cell transfer, mice were euthanized and subcutaneous tumors and spleens were harvested. Tumors were excised, weighed, and manually minced using sterile instruments. Visual inspection of tumor was performed such to avoid healthy adjacent tissue as well as normalize for both size and level of expected necrosis across all samples. After manual dissociation of tumors, enzymatic digestion was performed with collagenase D (400 U/mL), and DNase I (50 µg/mL) for 20 min at 37 °C with gentle agitation in RPMI 1640 medium supplemented with GlutaMAX™ (1x). Enzymatic digestion was terminated by the addition of 0.5 M EDTA (pH 8.0) at a final dilution of 1:100. Digested tumor samples were passed through 100-µm cell strainers, and cells were pelleted by centrifugation (600 × g, 5 min, 4 °C). Red blood cell lysis was performed using ACK lysis buffer for 1 min at room temperature. Cells were washed with PBS to obtain single-cell suspensions. Spleens were mechanically dissociated, filtered through 100-µm cell strainers, and centrifuged (600g, 5 min, 4 °C). Red blood cells were lysed using ACK buffer for 1 min at room temperature, followed by washing with PBS to generate single-cell suspensions.

**Tumor and spleen samples flow cytometry staining**

For T cell phenotyping studies, single-cell suspensions from tumors and spleens were stained 20 min in PBS supplemented with the following fluorochrome-conjugated antibodies: BV605 anti-mouse CD45 (clone 30-F11), PE-Cy7 anti-mouse CD3 (clone 17A2), BV510 anti-mouse CD8 (clone 53-6.7), Alexa Fluor™ 700 anti-mouse CD4 (clone RM4-5). T cell differentiation status was assessed using APC-Fire™ anti-mouse CD44 (clone IM7) and BV421 anti-mouse CD62L (clone MEL-14). T cell activation markers included PE anti-mouse CD137 (clone 17B5) and PerCP/Cy5.5 anti-mouse ICOS (clone 7E.17G9). T cell exhaustion markers were assessed using APC anti-mouse PD-1 (clone J43), BV785 anti-mouse TIM-3 (clone RMT3-23), and BV650 anti-mouse LAG-3 (clone C9B7W). CAR T cells were identified by GFP (FITC channel). For immune phenotyping studies the following fluorochrome-conjugated antibodies were used: BUV395 anti-mouse B220 (clone RA3-6B2), FITC anti-mouse Ly6G (clone 1A8), BV650 anti-mouse NKp46 (clone 29A1.4), PE-Cy7 anti-mouse CD11b (clone M1/70), BV785 anti-mouse PD-L1 (clone 10F.962), BV510 anti-mouse Ly6C (clone HK1.4), BV421 anti-mouse F4/80 (clone BM8), PE anti-mouse CD11c (clone N418), PerCP/Cy5.5 anti-mouse MHCII (I-A/I-E; clone M5/114.15.2), APC anti-mouse CD206 (clone C068C2), APC anti-mouse CD25 (clone PC61), BV421 anti-mouse TIGIT (clone 1G9), PE anti-mouse FoxP3 (clone FJK-16s), APC-H7 anti-mouse CD45 (clone 30-F11), and BV605 anti-mouse CD3 (clone 145-2C11). Cell viability was assessed using IR885 eosin-5′-maleimide (EMA) live/dead reagent Following staining, cells were washed, centrifuged (600g, 5 min, 4 °C), and resuspended in PBS supplemented with 0.05% BSA and 5 mM EDTA. Samples were acquired on a FACSCanto-II cytometer (BD Biosciences). Data were analyzed using FlowJo software (TreeStar).

**Tumor samples flow cytometry staining and sorting for scRNA-seq**

Tumor-derived single-cell suspensions were centrifuged (600g, 5 min, 4 °C) and incubated with anti-mouse CD16/32 Fc receptor-blocking antibody (Mouse Fc Block; Cat. No. 553142, BD Pharmingen) for 10 min at 4 °C. Following centrifugation, cells were then stained with TotalSeq™ hashing antibodies and surface markers for 30 min at 4 °C in PBS supplemented with 0.05% BSA and 5 mM EDTA at a concentration of $4 \times 10^6$ cells/mL. TotalSeq™ hashing antibodies enable sample multiplexing during single cell experiment [1] s. The following monoclonal anti-mouse MHC-I (clone M1/42) and anti-mouse CD45 (clone 30-F11) TotalSeq™ hashing antibodies (BioLegend) were used according to the manufacturer's recommendations: TotalSeq™-C0301 anti-mouse Hashtag 1 (Cat. No. 155861), TotalSeq™-C0302 anti-mouse Hashtag 2 (Cat. No. 155863), TotalSeq™-C0303 anti-mouse Hashtag 3 (Cat. No. 155865), and TotalSeq™-C0304 anti-mouse Hashtag 4 (Cat. No. 155867). Cell surface staining antibodies included PE anti-mouse CD45.2 (clone 104) and APC anti-mouse CD3 (clone 145-2C11). Cells were washed, centrifuged (450 × g, 5 min, 4 °C), and resuspended in PBS supplemented with 0.05% BSA and 5 mM EDTA. Cell viability was assessed using 7-aminoactinomycin D (7-AAD) at a 1:1,000 dilution immediately prior to FACS sorting. Immediately prior to sorting, samples were filtered through cell strainer caps and sorted on a BD FACSAria™ IIU cell sorter (BD Biosciences). $CD45^+$, $CD45^-$, and CAR-T ($GFP^+$) populations were collected for downstream single-cell RNA sequencing.

**scRNA-seq: Single cell data collection, cDNA amplification, library preparation and sequencing**

FACS-sorted cell populations were processed using the 10x Genomics Chromium platform for simultaneous single-cell transcriptome and T cell receptor (TCR) profiling. Four independent single-cell RNA-seq experiments were performed, two at day 4 and two at day 10 post-adoptive cell transfer. Each experiment included pooled tumor samples from PSMA-CAR-T- and EDA-CAR-T-treated mice (3-4 mice per experiment). From each mouse tumor sample, approximately 5,000 $CD45^+$ cells, 5,000 $CD45^-$ cells, and all recovered CAR-T cells ($GFP^+$) were included for library preparation.

**5′ gene expression profiling.** scRNA-seq was performed using the Chromium GEM-X Single Cell 5′ Reagent Kit v3 (10x Genomics) according to the manufacturer's instructions. Briefly, tumor cell suspensions were sorted in 100µl 1X PBS, 0.04% BSA and their concentration and viability were determined using Nexcelom's Cellometer K2 Fluorescent Cell Counter. Cell concentration was adjusted to ca. 2000 cells/µl, then mixed with RT mix and loaded in Chromium GEM-X single cell 5' Chips. Thus, individual cells were incorporated in Gel Beads in Emulsion (or GEMs), then lysed and the polyadenylated RNA was reverse-transcribed and barcoded. Barcoded cDNA was then recovered and PCR-amplified. Next, cDNA was quantified with Qubit dsDNA HS Assay Kit and its profile examined using Agilent's HS D5000 ScreenTape Assay. Fifty nanograms of double-stranded cDNA were used for 5′ Gene Expression library construction: cDNA was fragmented, end-repaired, A-tailed and ligated to an adaptor. A final PCR-amplification with barcoded primers allowed sample indexing. Sequencing was performed in a NextSeq2000 (Illumina) (Read1: 28 cycles; Read2: 90 cycles; i7 index: 10 cycles; i5: 10 cycles) at an average depth of 30,000-50,000 reads/cell.

**Single-cell TCR repertoire sequencing.** TCR α/β sequencing was performed with 10x Genomics Single Cell V(D)J Immune Profiling kit. Briefly, full-length V, D, and J gene segments were amplified from barcoded cDNA using the Chromium Single Cell V(D)J Enrichment Kit (for mouse T cells) via two consecutive PCRs with primers targeting both the adaptor and the gene constant region. Enriched cDNA was quantified as described above and 50 ng were further used for library construction as described above. After quality control and quantification, samples were pooled and sequenced in a NextSeq2000 (Illumina) (Read1: 28 cycles; Read2: 90 cycles; i7 index: 10 cycles; i5: 10 cycles) at an average depth of 7,000-15,000 reads per cell.

**Sample identification of cells through hashing.**
Sample identification was performed using 10x Cell Surface Protein Library Construction. In short, 5µl of small cDNA fragments were PCR amplified with indexed primers. Upon quality control, these libraries were sequenced in an Illumina NextSeq2000 (Read1: 28 cycles; Read2: ≥34 cycles; i7 index: 10 cycles; i5: 10 cycles) at an average depth of 150-500 reads per cell.

**Single-cell RNA-seq and V(D)J data processing**
Raw sequencing data were processed using Cell Ranger (v9.0.0, 10x Genomics) for demultiplexing, alignment to the mouse reference genome (GRCm38; refdata-gex-GRCm38-2020-A), barcode processing, and unique molecular identifier (UMI) counting. Downstream analysis of the gene-barcode matrix of UMI counts was performed using the Seurat R package (version 5.4.0, Satija Lab) which enables the integrated processing of multi-modal single cell datasets, including gene expression (RNA) and hashtag oligonucleotides (HTO). Ambient RNA contamination was estimated and removed using SoupX (v1.6.2). Both raw_feature_bc_matrix and filtered_feature_bc_matrix outputs from Cell Ranger were used as input. Potential doublets were identified using DoubletFinder (v2.0.4) (https://github.com/chris-mcginnis-ucsf/DoubletFinder), assuming a 10% expected doublet rate based on Poisson statistics. HTO-derived UMI counts were imported into Seurat as a separate HTO assay, normalized using centered log-ratio (CLR) transformation, and demultiplexed using the HTODemux function with a positive.quantile threshold of 0.95. Cells were classified as singlets, doublets, or negatives based on hashtag enrichment. Low-quality cells were excluded based on the following criteria: fewer than 400 detected genes, fewer than 50 or more than 40,000 total UMIs, mitochondrial transcript content exceeding 10%, or ribosomal transcript content exceeding 30%. Cells identified as doublets by either DoubletFinder or HTODemux were removed. Filtered count matrices were merged and log-normalized using the NormalizeData function. Highly variable features (n = 2,000) were identified using FindVariableFeatures, and data were scaled using ScaleData. Batch effects across experiments were corrected using Harmony with default parameters. T cell receptor (TCR) repertoire analysis was performed using the scRepertoire package.

**Clustering and cell type annotation**
Principal component analysis (PCA) was performed on scaled data, and the top 20 principal components were used for uniform manifold approximation and projection (UMAP), visualization, and unsupervised clustering. Clustering was performed at a resolution of 0.4. Major cell populations were annotated based on canonical marker gene expression profiles. Tumor cells derived from the F9 teratocarcinoma line were identified by expression of Zfp42, Bcat1, Bcat2, and Klf5. Fibroblasts were defined by enrichment of Dcn, Sparc, Pdpn, Pdgfra, Col1a1, and Col3a1. Immune populations were identified by Ptprc (CD45) expression and further subclassified as follows: T cells (Cd3e$^{+}$), including regulatory T cells (Tregs; Cd4$^{+}$ Foxp3$^{+}$); NK cells (Ncr1$^{+}$, Klrd1$^{+}$, Nkg7$^{+}$, Klrk1$^{+}$); granulocytes (S100a8$^{+}$, S100a9$^{+}$, Ccr1$^{+}$, Cxcr2$^{+}$); monocytes/macrophages (Csf1r$^{+}$, Ly6c2$^{+}$, Cd14$^{+}$, Cd68$^{+}$, Adgre1$^{+}$, H2-Aa$^{+}$); type 1

conventional dendritic cells (cDC1; H2-Aa$^+$, Cd80$^+$, Cd86$^+$, Flt3$^+$, Ccr7$^+$); plasmacytoid dendritic cells (pDCs; H2-Aa$^+$, Flt3$^+$, Bst2$^+$, Siglech$^+$, Clec10a$^+$, Clec12a$^+$); and basophils (Mcpt8$^+$, Gata2$^+$, Cd200r3$^+$). Module scores for predefined gene signatures were computed using the AddModuleScore function.

For higher-resolution analyses, T cells (including Tregs) and myeloid populations were subsetted, reprocessed, and reclustered using the top 20 principal components, with clustering resolutions of 0.6 for T cells and 0.5 for myeloid cells. Cluster-specific marker genes were identified using FindAllMarkers (average log fold change > 0.25; detected in >10% of cells). Marker expression patterns were visualized using dot plots and feature plots, and final subtype annotation was based on cluster-specific transcriptional profiles combined with established canonical markers. Within the T cell compartment, memory-like clusters were defined by coordinated expression of Tcf7, Ccr7, Sell, Cd27, and Il7r. Effector T cell clusters were characterized by high Cd44 expression together with cytotoxic and effector-associated genes including Prf1, Gzma, Gzmb, Gzmk, Ifng, and Cd40lg. These effector states frequently co-expressed activation and costimulatory molecules (Cd28, Tnfrsf4, Tnfrsf9, Tnfrsf18, Il2ra, Icos) alongside inhibitory and exhaustion-associated genes (Ctla4, Pdcd1, Lag3, Havcr2, Tox, Tigit). Intermediate effector-memory populations displayed mixed transcriptional programs, combining memory- and effector-associated features, indicative of transitional differentiation states. Proliferating T cells were identified by elevated expression of canonical cell-cycle genes, including Mki67, Tuba1b, Rrm1, and Smc2. In addition to these principal subsets, smaller populations were resolved, including Tregs (Cd4$^+$ Foxp3$^+$ Il2ra$^+$ Icos$^+$ Tigit$^+$ Il10$^+$), Th17-like CD4$^+$ T cells (Rorc$^+$ Il17ra$^+$), interferon-responsive CD8$^+$ T cells (Isg15$^+$ Isg20$^+$ Irf7$^+$), and cytotoxic γδ T cells (Trgv$^+$ Trdv$^+$ Prf1$^+$ Gzmb$^+$). CAR-T cells were identified and subset from the T cell subclusters based on GFP expression. Cells with at least one detected GFP count were annotated as CAR-T cells.

Within the immune compartment, macrophages were defined by expression of Csf1r, H2-Aa, Cd80, Cd83, and Cd86, whereas monocytes were characterized by Ly6c2 and Csf1r expression. Granulocytes segregated into three transcriptionally distinct subsets based on chemokine receptor expression and interferon-response signatures. A subset with higher expression of Cxcr2 and Cxcr4 was classified as resident granulocytes, whereas cells with comparatively lower expression of these receptors were designated as circulating granulocytes. A third population, characterized by strong enrichment of interferon-stimulated genes including Rsad2, Isg15, and Irf7, was defined as the IFN-response granulocyte subset. Macrophages also resolved into three transcriptionally distinct subsets reflecting functional specialization. An antigen-presentation subset was defined by high expression of MHC class II genes, including H2-Aa. A second subset, termed the endothelial-migratory macrophages, was enriched for genes associated with endothelial interaction, adhesion, and migration, such as Pecam1, Itga4, Itgal, Pparg, and Ace. Finally, an immunosuppressive macrophage subset displayed elevated expression of canonical M2-like markers, including Mrc1 and Sirpa, consistent with regulatory and suppressive functions within the tumor microenvironment.

**Pseudobulk differential gene expression and enrichment analyses**

Differential gene expression (DGE) analyses of CAR-T cells, T cells and myeloid populations were performed using a pseudobulk strategy. For each defined cell population, raw UMI counts were aggregated at the level of biological sample and analyzed using DESeq2, which models count data with a negative binomial generalized linear model. Condition-dependent transcriptional changes were assessed using a likelihood ratio test (LRT), comparing a full model including treatment condition to a reduced intercept-only model, with PSMACAR-T set as the reference condition. Automatic outlier replacement was disabled to preserve biologically meaningful variability. P-values were adjusted for multiple testing using the Benjamini-Hochberg false discovery rate (FDR), and genes with FDR < 0.05 and an absolute $\log_2$ fold change above threshold were considered differentially expressed.

Functional pathway analyses were conducted using the clusterProfiler package. For Gene Ontology (GO) enrichment analysis, genes from pseudobulk DESeq2 results were filtered based on absolute log2 fold change and nominal P-value < 0.2. Enrichment of GO Biological Process terms was assessed using the enrichGO function with the org.Mm.eg.db annotation database. Statistical significance was evaluated using a hypergeometric test with a P-value cutoff of 0.01 and an FDR threshold of 0.05.

Gene set enrichment analysis (GSEA) was conducted using ranked gene lists based on signed $\log_2$ fold changes from DESeq2. Curated gene signatures related to T cell activation, cytotoxicity, exhaustion, and immune function were used. Pathways with an FDR < 0.25 were considered significantly enriched, in accordance with established GSEA guidelines.

**Gene set variation analysis and definition of T cell scores and gene sets**

Gene set variation analysis (GSVA) was performed using the GSVA R package, applying a non-parametric, unsupervised approach to assess pathway-level activity across conditions. Custom gene sets were curated to capture functional programs most relevant to CAR-T and T cell biology within the TME. These included signatures related to cytotoxic effector function, terminal exhaustion, T cell activation and helper function. Gene sets were defined based on a combination of previously published signatures, canonical immune effector genes, and markers consistently identified in our single-cell differential expression and GSEA enrichment analyses.
GSVA scores were calculated from log-transformed pseudobulk expression matrices. Differences across conditions were evaluated using the Kruskal-Wallis test followed by Dunn's post hoc test with Benjamini-Hochberg correction. Pairwise comparisons were performed using the Wilcoxon rank-sum test where indicated. Adjusted P-values < 0.05 were considered statistically significant.

**Cell-cell communication analysis**

CellChat was used for analyzing intercellular communication networks based on the receptor-ligand interactions from multiple databases. The scRNA-seq input data for CellChat included quantitative count data and cell-type annotation information. In brief, the percentage and the average of gene expressing for each gene in the cells were calculated. The ligand-receptor pairs were filtered to obtain receptor and ligand genes exceeding a specified threshold (the default is 10%). A pair-to-pair comparison was then performed between all cell types in the dataset, and the actual mean value of the ligand-receptor pairs between two cell types was calculated to speculate p-value of the receptor-ligand pair in 2 cell types. Finally, the highly specific interactions between cell types were arranged through the enrichment results of significant ligand receptor pairs.

**Statistical analysis**

Bioinformatic and statistical analyses of single-cell RNA-seq data were conducted using R software (version 4.2.2; R Foundation for Statistical Computing, Vienna, Austria; https://www.r-project.org) and RStudio, using the packages described above. Single-cell data processing and downstream analyses included Louvain clustering and cell-cell communication inference using CellChat. For cell-cell communication analyses using CellChat, interaction strengths were standardized as Z-scores relative to background distributions, with positive and negative values indicating increased or decreased signaling activity, respectively. Z-scores were used as effect-size measures, and statistical significance was assessed using the permutation-based or model-based frameworks implemented in CellChat. Unless otherwise specified, multiple testing correction was performed using the Benjamini-Hochberg false discovery rate (FDR), with adjusted P-values < 0.05 considered statistically significant. Absolute Z-scores > 2 were used as a predefined threshold to highlight biologically relevant differences in signaling activity. Conventional statistical analyses were performed using GraphPad Prism version 7 (GraphPad Software). Parametric tests included one-way analysis of variance (ANOVA) followed by Bonferroni's multiple-comparison correction and two-way ANOVA followed by Tukey's multiple-comparison correction. Non-parametric tests included tthe Kruskal-Wallis test. Statistical analyses were performed on biological replicates, as indicated in the figure legends. For all tests, P-values < 0.05 were considered statistically significant. Unless otherwise stated, data are presented as mean ± standard deviation (SD). P-values < 0.05 were considered statistically significant. Statistical significance is denoted as *P < 0.05, **P < 0.01, ***P < 0.001, and ****P < 0.0001.

### SUPPLEMENTARY FIGURES

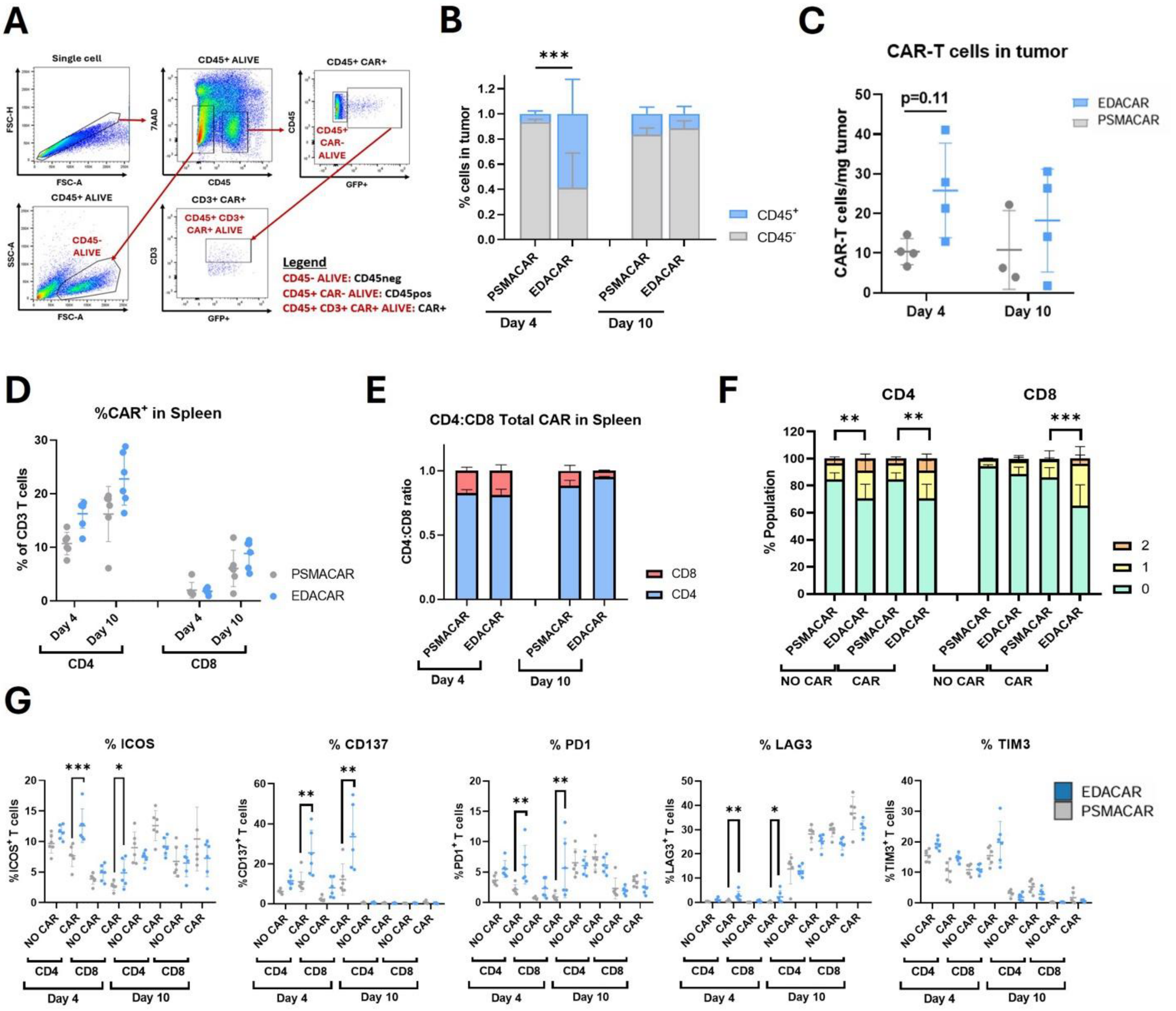


**Fig. S1. Flow cytometry gating strategy for sorting tumor-infiltrating immune populations for scRNA-seq and quantification of splenic T cell and CAR-T cell populations. (A)** Flow cytometry gating strategy used to isolate CD45$^{-}$, CD45$^{+}$, and CAR-T (GFP$^{+}$ CD3$^{+}$) cell populations from dissociated tumor samples for scRNA-seq. **(B)** Ratio of CD45$^{+}$ to CD45$^{-}$ cells in tumors from αPSMA- or EDACAR-T-treated mice at day 4 and day 10 post-infusion. **(C)** Quantification of number of CAR-T cells per mg of tumor tissue at day 4 and day 10 post-infusion for αEDA- and PSMACAR-T treatment groups. **(D)** Percentage of CAR-T cells among CD3$^{+}$ T cells in the spleen from αPSMA- or EDACAR-T-treated mice at day 4 and day 10 post-infusion. **(E)** CD4:CD8 ratio of CAR-T cells in the spleen from αPSMA- or EDACAR-T-treated mice at day 4 and day 10 post-infusion. **(F)** T cell activation of T cells and CAR-T cells in the spleen from αPSMA- or EDACAR-T-treated mice at day 4 post-infusion. T cell activation markers: ICOS, CD137. **(G)** Percentage of ICOS, CD137, PD1, LAG3 and TIM3 positive T cells and CAR-T cells in the spleen from αPSMA- or EDACAR-T-treated mice at day 4 and day 10 post-infusion. B-E, day 4 (n:4 PSMACAR-T and EDACAR-T biological replicates) and day 10 (n:3 PSMACAR-T and n:4 EDACAR-T biological replicates); F-G, day 4 and day 10 (n:6 PSMACAR-T and EDACAR-T biological replicates). Two-way ANOVA with Tukey's multiple comparisons test (B, C, F and G). Bars and lines representing the mean and SD are plotted (B-G). Statistical significance is denoted as *P < 0.05, **P < 0.01 and ***P < 0.001. ANOVA, analysis of variance.

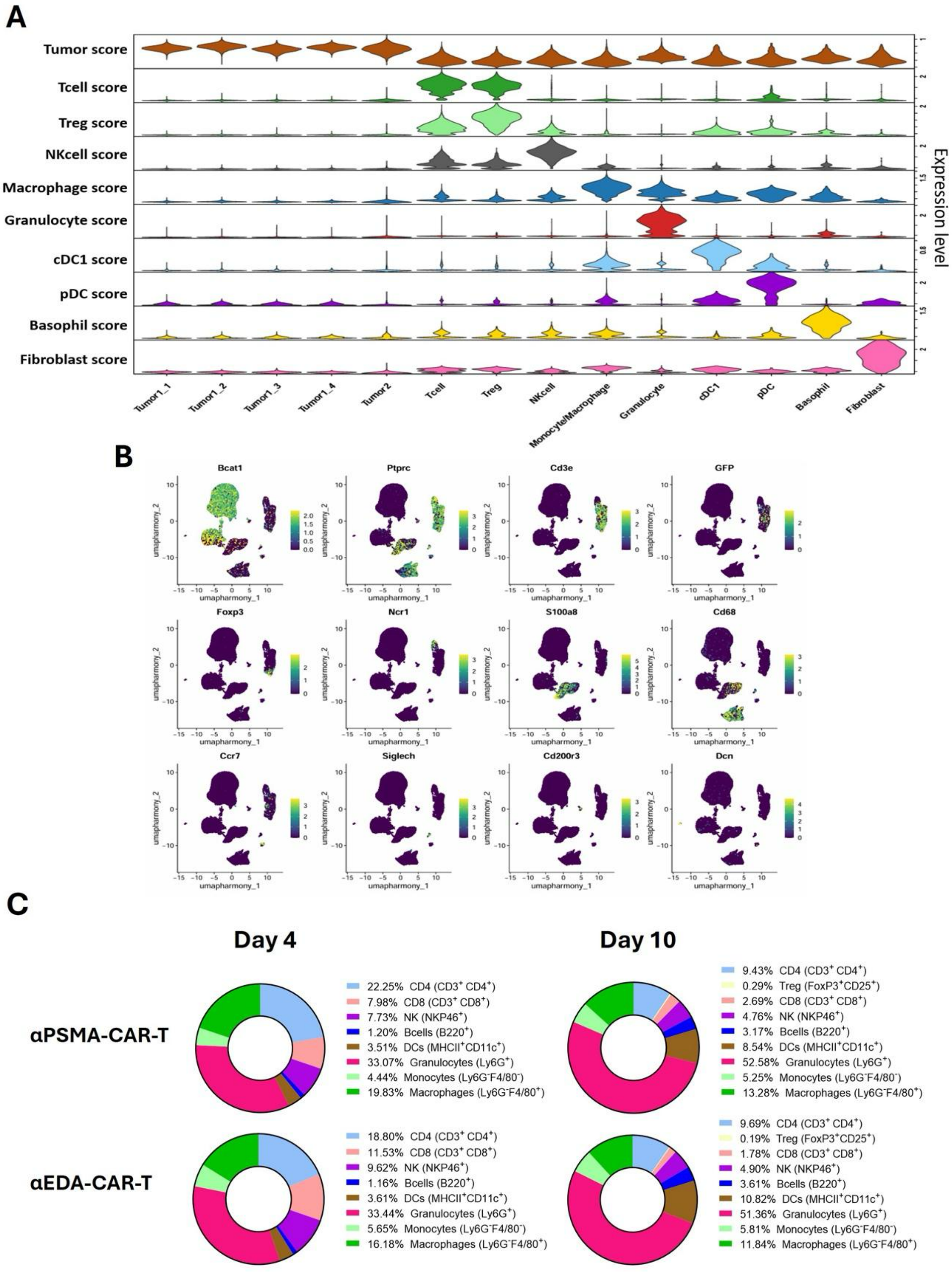


**Fig. S2. Identification and annotation of major TME populations. (A)** Distribution of curated lineage signature scores across major TME populations identified. **(B)** Feature plots displaying the relative expression of representative marker genes used to annotate the main TME populations. **(C)** Immune phenotyping of PSMACAR-T- or EDACAR-T-treated tumors at day 4 and day 10 post-infusion by flow cytometry. Day 4 and day 10 (n:4-5 PSMACAR-T and EDACAR-T biological replicates).

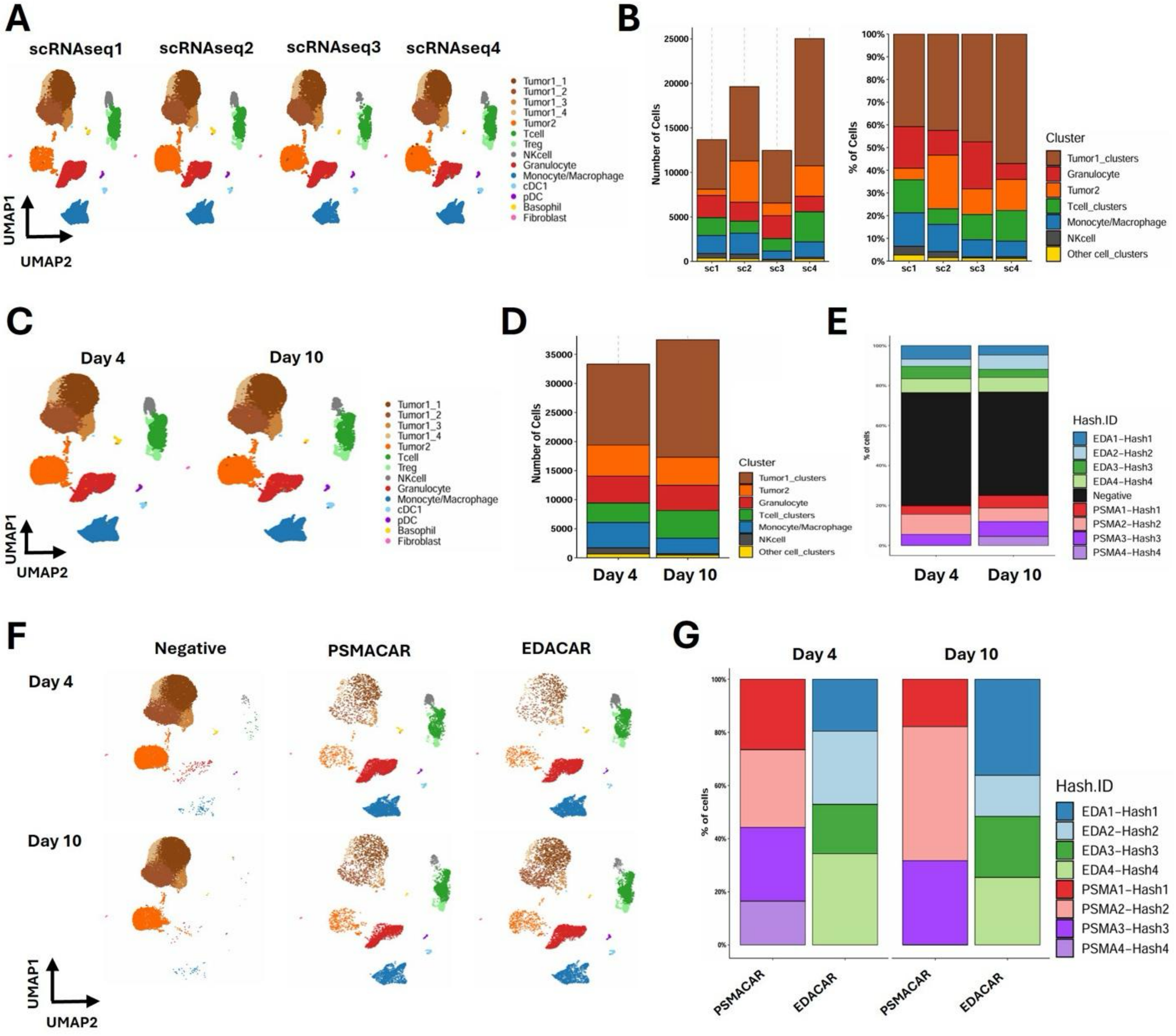


**Fig. S3. Quality control of the scRNAseq dataset. (A)** UMAP plots showing the distribution of major TME populations across the four independent scRNA-seq experiments. Datasets sc1-sc2 correspond to day 4 post-infusion and sc3-sc4 to day 10 post-infusion. Each experiment contains tumor samples treated with either PSMACAR-T or EDACAR-T cells. **(B)** Absolute cell numbers (left) and proportional representation (right) of annotated TME populations across independent scRNA-seq experiments. **(C)** UMAP plots showing TME population distributions grouped by time point (day 4 vs. day 10 post-infusion). **(D)** Absolute number of cells per TME population across day 4 and day 10 post-infusion samples. **(E)** Distribution of hashing identities across biological replicates at each time point (day 4 and day 10 post-infusion). **(F)** UMAP visualization of TME populations split by hashing identity across day 4 and day 10 post-infusion. Negative indicates cells for which no hashing-based identity could be resolved. **(G)** Contribution of treatment-defined hashing identities (PSMACAR-T vs. EDACAR-T) across biological replicates at day 4 and day 10 post-infusion.

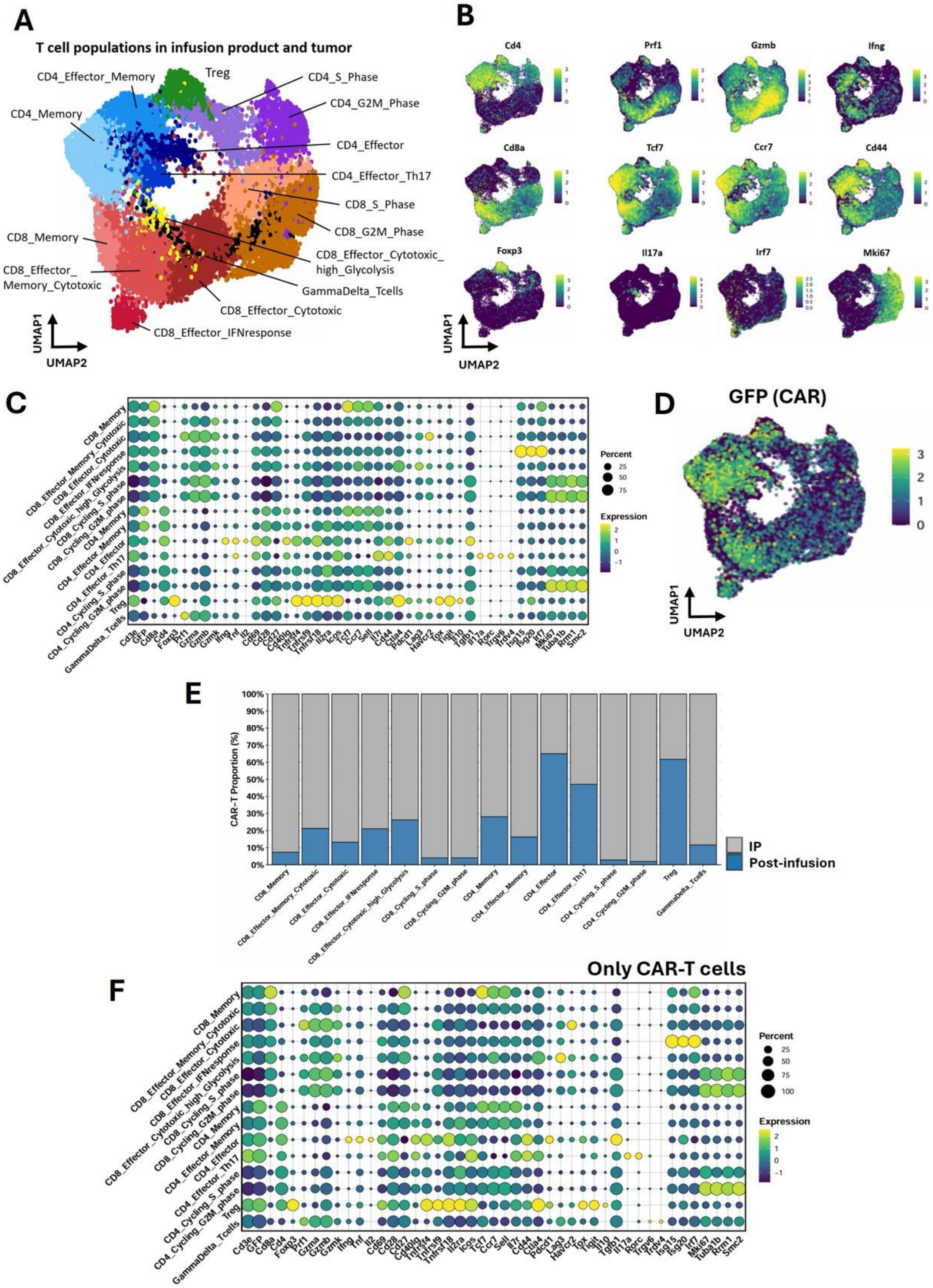


**Fig. S4. Identification and annotation of T and CAR-T cell populations in the TME. (A)** UMAP plot showing T cell subclusters identified across the infusion product and post-infusion tumor samples. **(B)** Feature plots displaying key identity markers for each T cell subcluster. **(C)** Average expression matrix of major marker genes used to define T cell subclusters. **(D)** Relative expression of Gfp (GFP, CAR marker) across T cell subclusters. **(E)** Proportion of CAR-T cells across T cell subclusters in the infusion product (IP) and the post-infusion tumor samples. **(F)** Average expression matrix of major marker genes used to define CAR-T-cell subclusters.

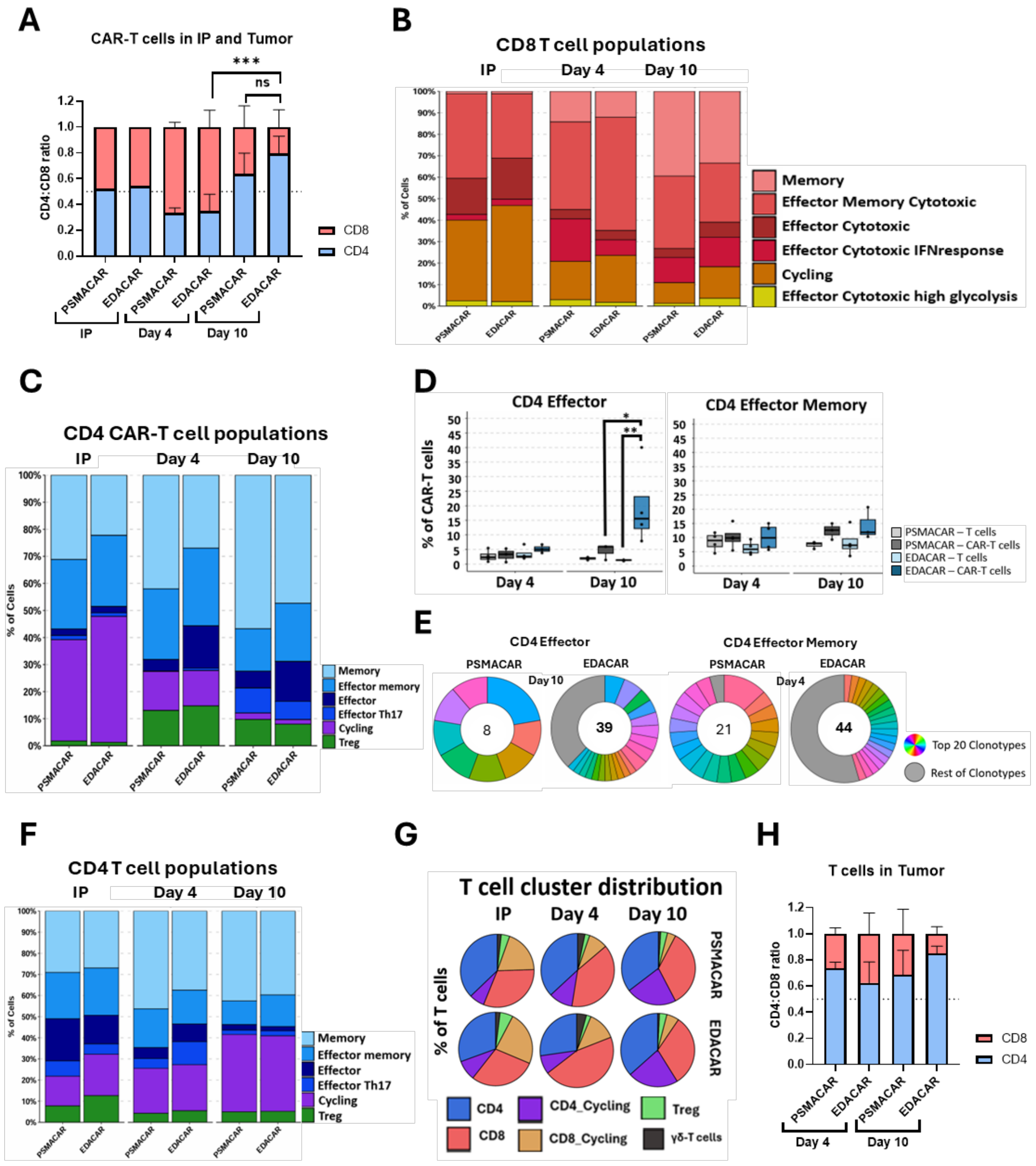


**Fig. S5. CD4 and CD8 characterization in the TME. (A)** CD4:CD8 ratio of CAR-T cells in the infusion product (IP) and tumor from PSMACAR-T- or EDACAR-T-treated mice at day 4 (n:6 PSMACAR-T and EDACAR-T biological replicates) and day 10 (n:4 PSMACAR-T and n:5 EDACAR-T biological replicates) post-infusion by flow cytometry. Bars representing the mean and SD are plotted. **(B)** Distribution of $CD8^{+}$ T cell subclusters across infusion product and post-infusion tumor samples. **(C)** Distribution of $CD4^{+}$ CAR-T cell subclusters across the infusion product (IP) and post-infusion tumor samples (day 4 and day 10). **(D)** Abundance of $CD4^{+}$ effector and effector-memory T-cell and CAR-T cell subsets across biological replicates at day 4 and day 10 post-infusion. The box represents the interquartile range, the line indicates the median, and whiskers denote minimum to maximum values. **(E)** Clonotype diversity of $CD4^{+}$ effector CAR-T cells at day 10 post-infusion and $CD4^{+}$ effector-memory CAR-T cells at day 4 post-infusion. **(F)** Distribution of CD4$^+$ T cell subclusters across infusion product and post-infusion tumor samples. **(G)** Composition of major T cell subsets in the infusion product (IP) and at day 4 and day 10 post-infusion tumor samples. **(H)** CD4:CD8 ratio of T cells in the tumor from PSMACAR-T- or EDACAR-T-treated mice at day 4 (n:6 PSMACAR-T and EDACAR-T biological replicates) and day 10 (n:4 PSMACAR-T and n:5 EDACAR-T biological replicates) post-infusion by flow cytometry. Two-way ANOVA with Tukey's multiple comparisons test. Bars representing the mean and SD are plotted. Two-way ANOVA with Tukey's multiple comparisons test (A and C). Statistical significance is denoted as *Padj < 0.05, **Padj < 0.01 and ***P < 0.001. ANOVA, analysis of variance.

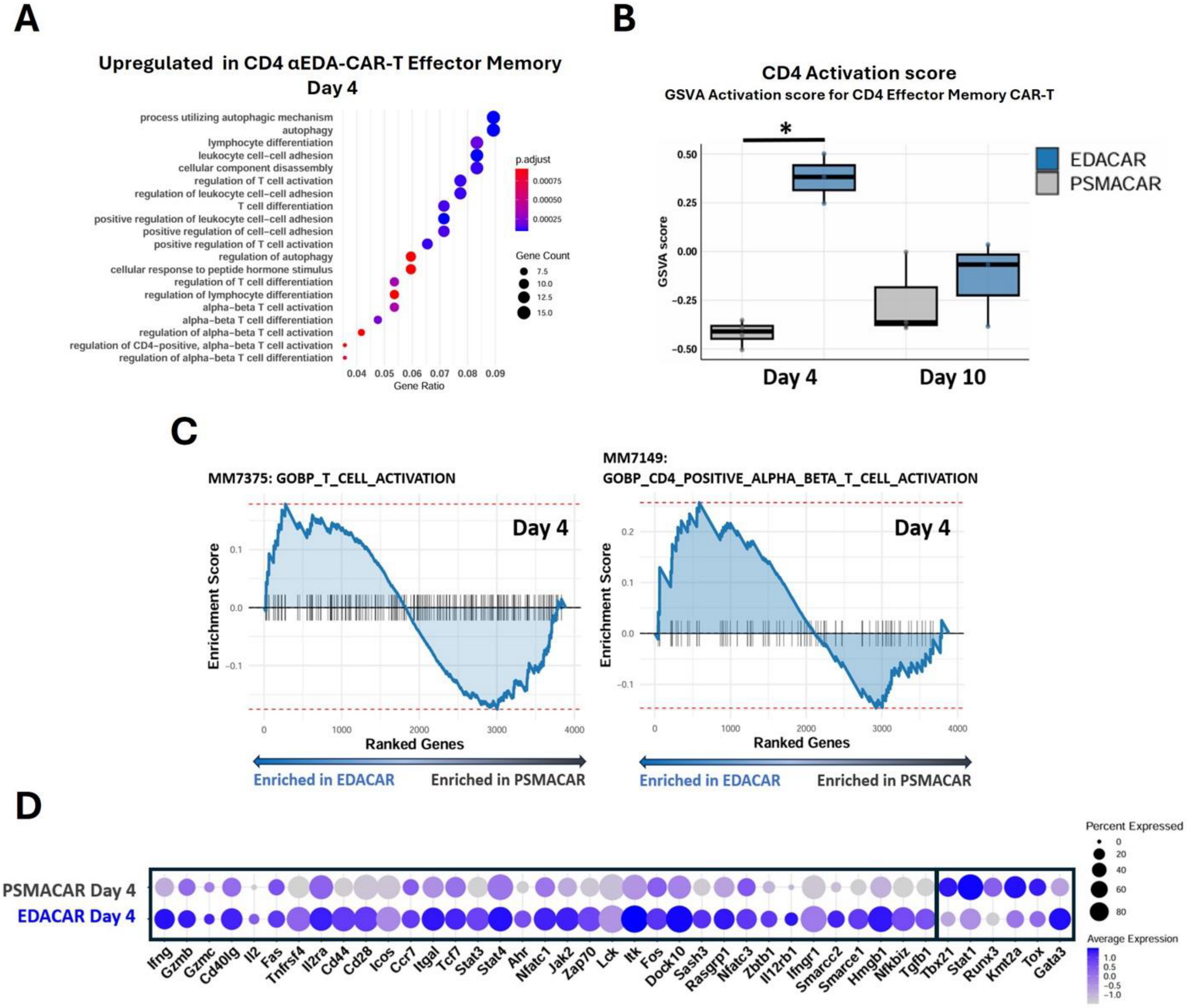


**Fig. S6. Transcriptional reprogramming of effector memory CD4⁺ EDACAR-T cells. (A)** Differential Gene Ontology enrichment analysis of genes upregulated in effector memory CD4⁺ EDACAR-T cells compared with PSMACAR-T cells at day 4 post-infusion. **(B)** GSVA scoring of effector memory CD4⁺ CAR-T cells. Pseudobulk GSVA scoring of effector memory CD4⁺ CAR-T cells at day 4 using a custom *CD4 activation* gene set. Kruskal-Wallis test followed by Dunn's post hoc test with Benjamini-Hochberg correction. Pairwise comparisons were performed using the Wilcoxon rank-sum test. The box represents the interquartile range, the line indicates the median, and whiskers denote minimum to maximum values. **(C)** GSEA of curated CD4⁺ T-cell activation signatures (MM7375, MM7149) comparing effector memory CD4⁺ EDACAR-T cells and control PSMACAR-T cells at day 4 post-infusion. **(D)** Average expression of representative genes included in the custom CD4 Activation gene set used for pseudobulk. Statistical significance is denoted as *Padj < 0.05. GO, gene ontology; GOBP; gene ontology biological processes; GSVA, Gene Set Variation Analysis.

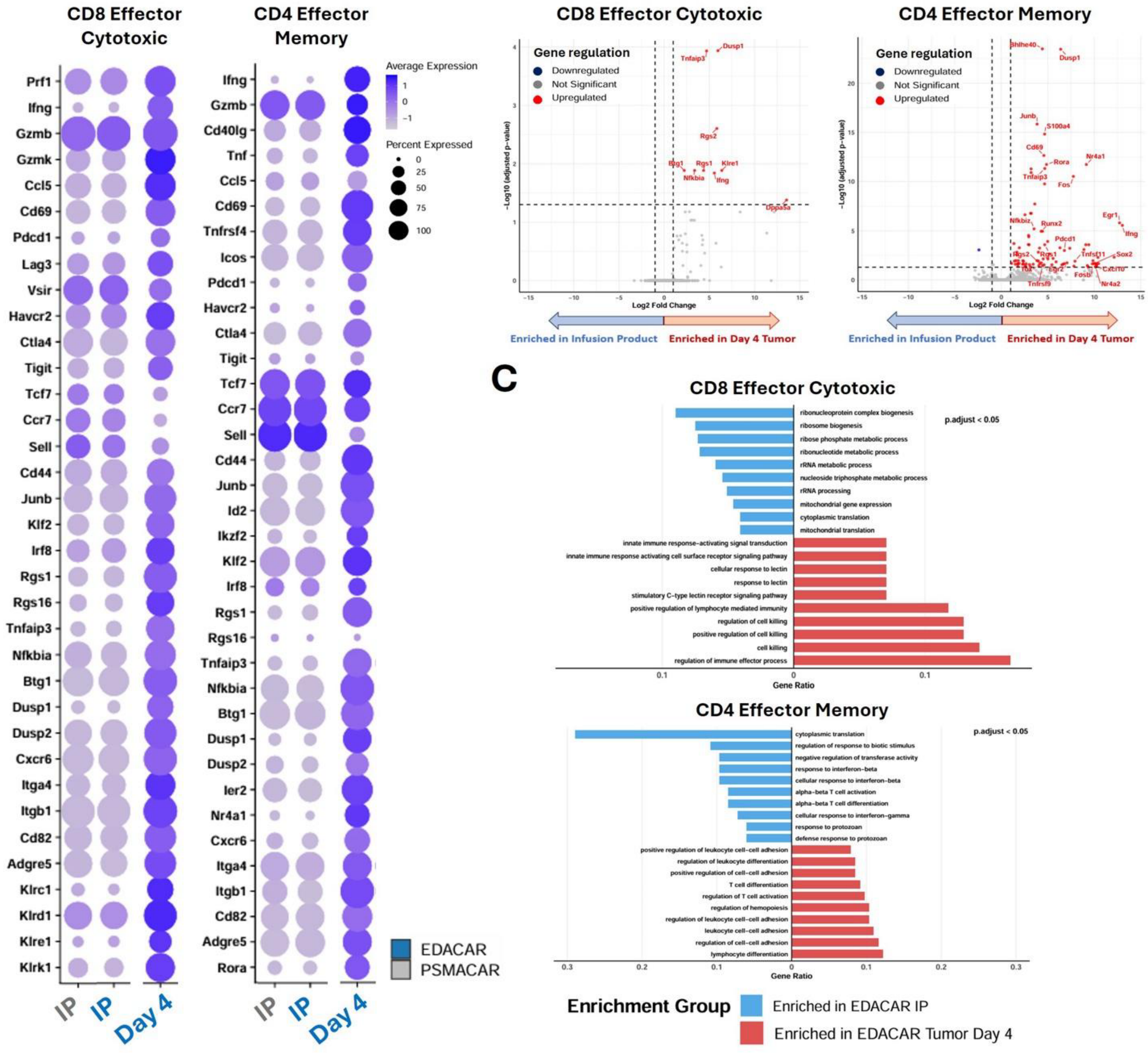


**Fig. S7. Transcriptional rewiring of EDACAR-T cells from the infusion product to the tumor. (A)** Average expression of key T-cell identity, activation, and effector-function markers in effector cytotoxic CD8⁺ (left) and effector memory CD4⁺ (rigth) EDACAR-T cells from the infusion product (IP; blue), compared with PSMACAR-T cells from the infusion product (grey), and EDACAR-T cells isolated from tumors at day 4 post-infusion (Day 4; blue). **(B)** Differentially expressed genes in effector cytotoxic CD8⁺ (left) and effector memory CD4⁺ (rigth) EDACAR-T cells recovered from tumors at day 4 post-infusion (red), compared with matched EDACAR-T cells from the infusion product (blue). **(C)** Differential Gene Ontology enrichment analysis of biological processes associated with genes upregulated in EDACAR-T cells at day 4 post-infusion (red) or enriched in the infusion product (blue), shown separately for effector cytotoxic CD8⁺ (up) and effector memory CD4⁺ CAR-T subsets (down).

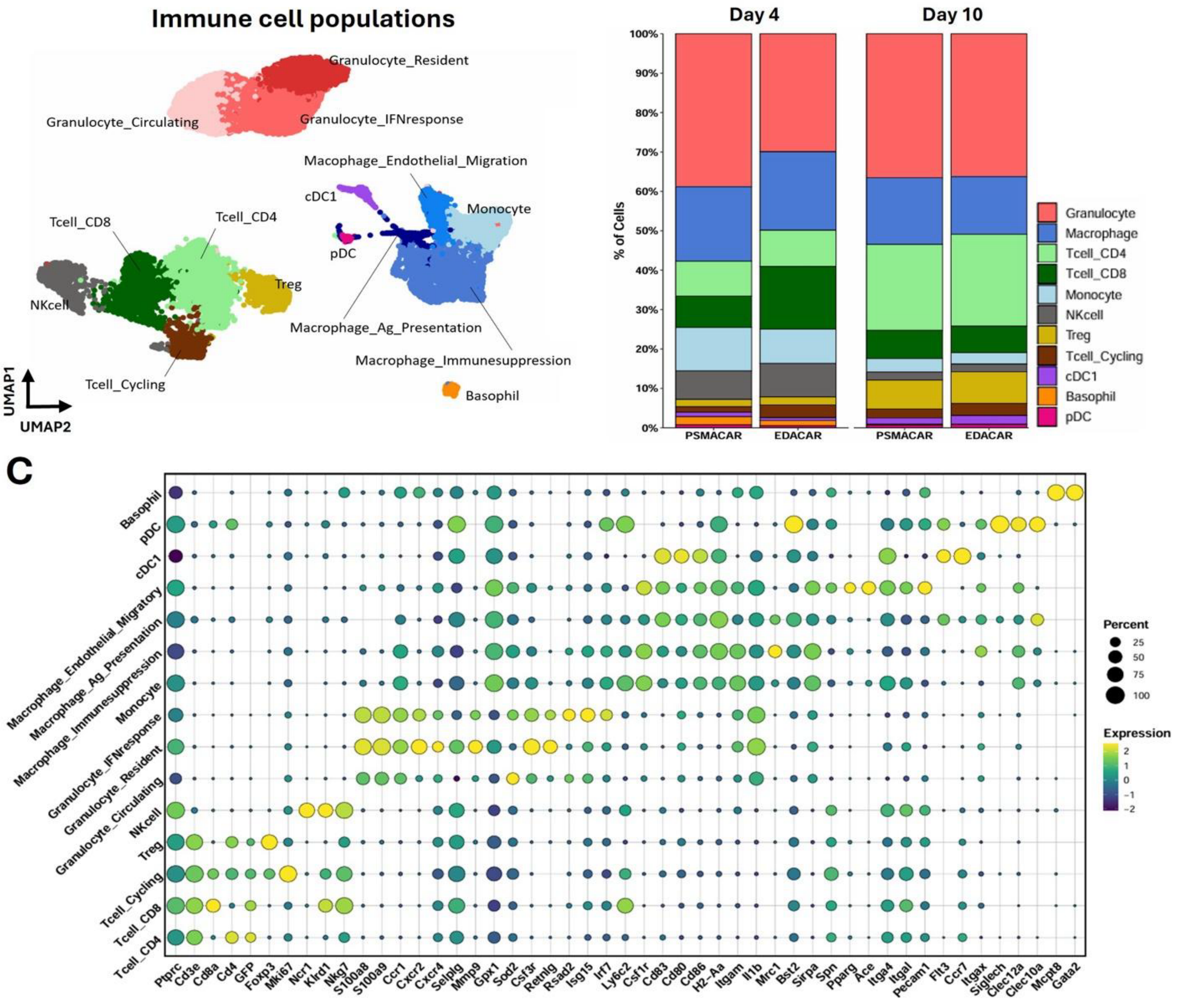


**Fig. S8. Identification and annotation of immune cell populations. (A)** UMAP visualization of immune cell subclusters identified across tumor samples. **(B)** Proportion of major immune cell types in PSMACAR-T- and EDACAR-T-treated tumors at day 4 and day 10 post-infusion. **(C)** Average expression matrix of canonical identity markers across immune cell subclusters.

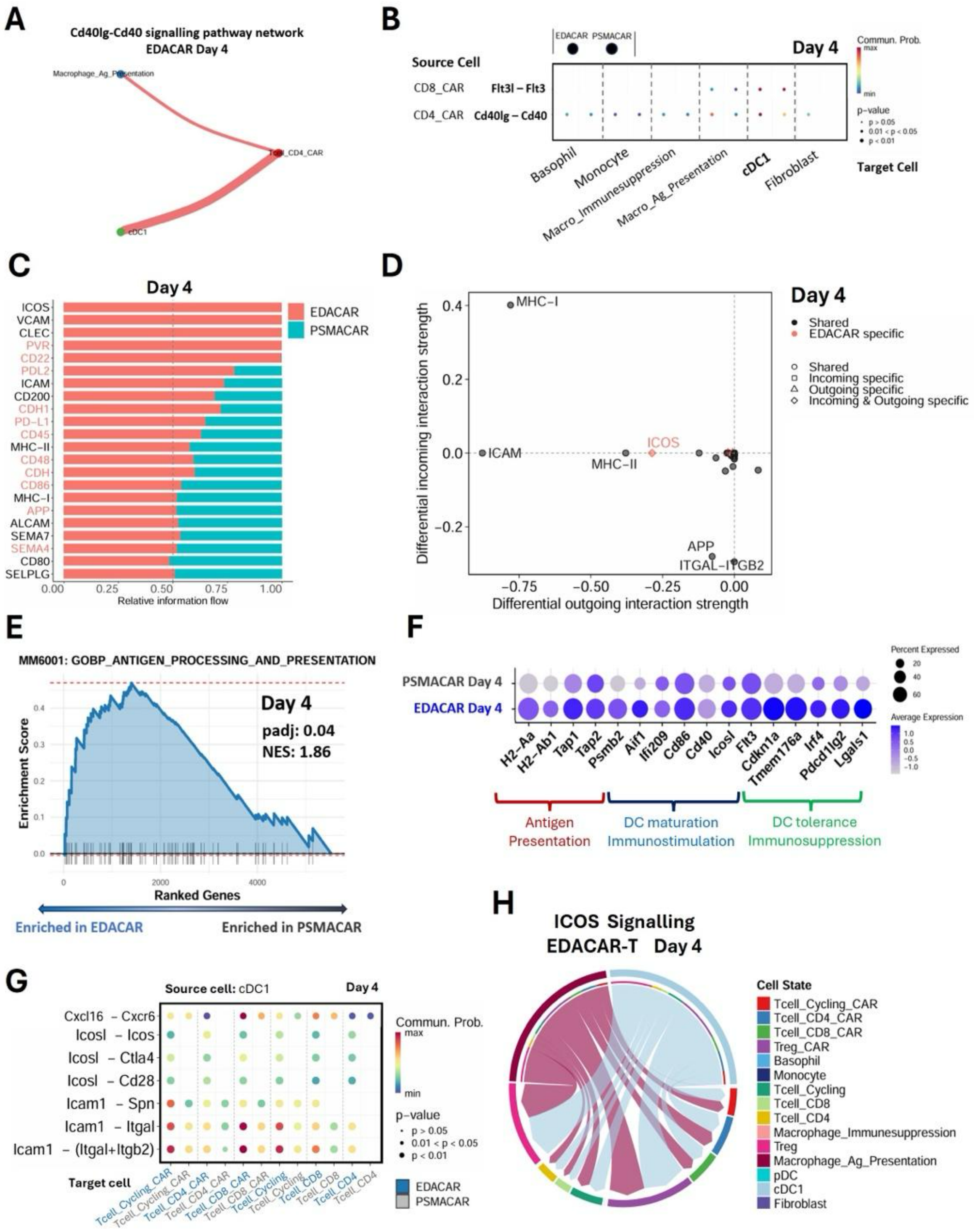

**Fig. S9. cDC1 communication networks with CAR-T cells. (A)** Circle plot showing the strength of CD40LG-CD40 interactions in the TME from EDACAR-T cells (source) at day 4 post-infusion. **(B)** Differential CD40 and FLT3L ligand-receptor interactions from CD4⁺ and CD8⁺ EDACAR-T cells (source) compared with control PSMACAR-T cells at day 4 post-infusion. For each target cell population, the left dot represents the interaction probability in EDA-CAR-T-treated tumors, whereas the right dot represents the interaction probability in PSMA-CAR-T-treated control tumors. **(C)** Differential overall information flow within the inferred Cell-Cell Contact signaling network for cDC1 (source) in EDACAR-T-treated tumors relative to PSMACAR-T-treated tumors at day 4 post-infusion. **(D)** Differential incoming and outgoing signaling strength within the inferred Cell-Cell Contact network for cDC1 (source) from EDACAR-T-treated tumors compared to control PSMACAR-T-treated at day 4 post-infusion. **(E)** GSEA analysis showing differential enrichment of the antigen processing and presentation gene signature (MM6001) in cDC1 from EDACAR-T-treated tumors compared with PSMACAR-T-treated tumors at day 4 post-infusion. **(F)** Average expression of cDC1 marker genes associated with antigen presentation, dendritic cell maturation, immunostimulation, and immunoregulatory/tolerogenic functions at day 4 post-infusion. **(G)** Differential CXCL16, ICOS, and ICAM1 ligand-receptor interactions between cDC1 (source) and T cells or CAR-T cells (target) from EDACAR-T-treated tumors compared to control PSMACAR-T-treated at day 4 post-infusion. **(H)** Chord plot showing the strength of ICOS interactions across the TME in EDACAR-T-treated tumor at day 4 post-infusion.

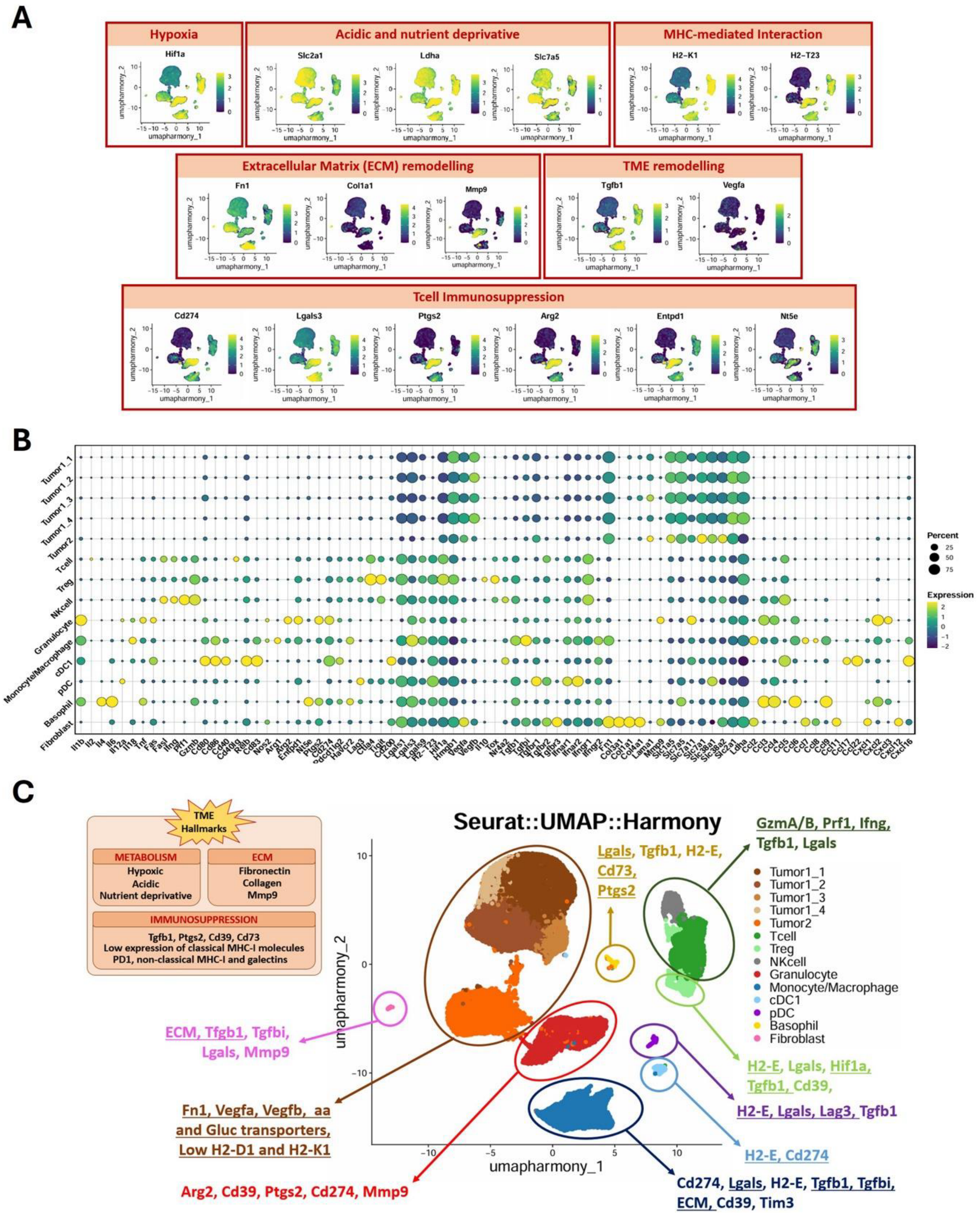


**Fig. S10. Identification of immunosuppressive and metabolically hostile TME hallmarks. (A)** Feature plots showing the expression of hallmark gene sets associated with major immunosuppressive and hostile TME programs. **(B)** Average expression matrix of key genes associated with immunosuppressive and metabolically hostile TME hallmarks across major TME populations. **(C)** Schematic summary outlining the dominant immunosuppressive and metabolically restrictive TME features inferred from single-cell transcriptomic analyses, highlighting population-specific contributions to hypoxia, ECM remodelling, immunoregulatory ligand expression, and T cell-suppressive pathways.

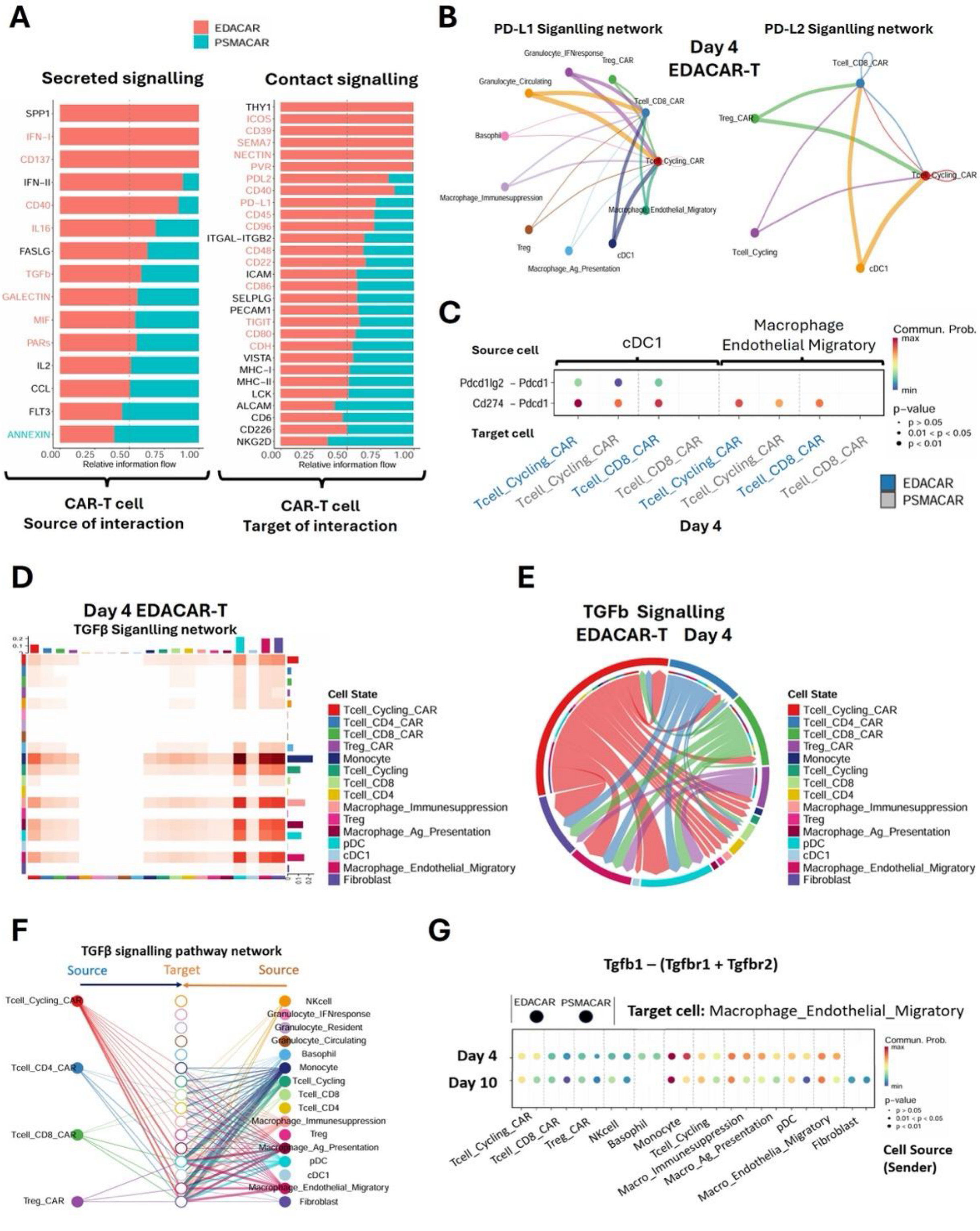


**Fig. S11. EDACAR-T cells engage PD-L1, PD-L2 and TGFβ immunosuppressive signaling networks within the TME. (A)** Differential overall information flow in the inferred Secreted Signalling (left) and Cell-Cell Contact (right) networks between EDACAR-T and PSMACAR-T cells at day 4 post-infusion. Left: CAR-T cells as signal senders. Right: CAR-T cells as signal receivers. **(B)** Circle plot showing the strength of PD-L1 and PD-L2 signalling interactions to EDACAR-T cells (target) in the TME at day 4 post-infusion. **(C)** Differential PD-L1 and PD-L2 ligand-receptor interactions between cDC1 and Macrophages_Endothelial_Migratory (source) and T cells and CAR-T cells (target) in EDACAR-T-treated tumors compared with PSMACAR-T controls at day 4 post-infusion. **(D)** Inferred TGFβ signaling interactions among TME populations at day 4 post-infusion. **(E)** Chord plot showing the global TGFβ signaling strength landscape in EDACAR-T-treated tumors at day 4 post-infusion. **(F)** Hierarchical representation of TGFβ pathway interactions between CAR-T cells and TME populations at day 4 post-infusion. Filled nodes represent signalling sources; open nodes represent targets. Edge colors correspond to the source cell type. **(G)** Differential TGFβ ligand-receptor interactions towards Macrophages_Endothelial_Migratory (target) in the TME from EDACAR-T-treated tumors compared to control PSMACAR-T-treated at day 4 and 10 post-infusion. For each cell source population, the left dot represents the interaction probability in EDA-CAR-T-treated tumors, whereas the right dot represents the interaction probability in PSMA-CAR-T-treated control tumors.

### SUPPLEMENTARY TABLES

**Supplementary Table S1.** Summary of cell loading, quality control metrics, and experimental design summary of the scRNA-seq experiments.

**Supplementary Table S2.** Total number of cells sorted from tumor samples at day 4 and day 10 post-infusion.

**Supplementary Table S3.** Gene sets used for defining transcriptional signatures and annotation of major TME cell populations.

**Supplementary Table S4.** Statistical results of GSEA.

**Supplementary Table S5.** Statistical results of GO enrichment analysis.

**Supplementary Table S6.** Custom gene sets used for T cells and GSVA enrichment results.

**Supplementary Table S7.** Statistical results of DGE analysis of the IP.

**Supplementary Table S8.** Statistical results of cell-cell communication analysis inferred by CellChat.
